# Optogenetic Rescue Reveals Spatiotemporal Rules of Germ-Layer Patterning

**DOI:** 10.64898/2025.12.08.693069

**Authors:** Naomi Baxter, Robert Piscopio, Joseph Rufo, Dasol Han, Isobel Whitehead, Jasmine Dhillon, Siddharth S. Dey, Maxwell Z. Wilson

## Abstract

Embryonic cells must interpret morphogen signals that vary in both time and space, but the rules by which they decode these dynamics remain unclear. Here we combine optogenetics with human 2D gastruloids to define minimal WNT signaling rules for germ-layer patterning. We block endogenous WNT secretion to create a “blank canvas” and reconstitute signaling using light-gated LRP6. Systematic temporal scans reveal a narrow competence window when the onset and duration of WNT signaling specify mesoderm; this window is shifted by cell density and amplified by BMP priming, whereas identical WNT inputs outside it invert germ-layer order or generate alternative mesodermal subtypes. Using micromirror-based illumination, we restricted WNT activation to a mid-ring during this temporal window; combined with BMP4, this fully restored germ layer domains with boundaries sharper than those generated by ligand stimulation. Thus, precise spatiotemporal control of a single pathway is sufficient to optically rebuild germ-layer architecture and reveals WNT as a temporal morphogen.

## Introduction

A central aim of developmental biology is to define the minimal requirements for tissue patterning. Classic organizer experiments demonstrated that certain signals are sufficient to induce new axes^1^, but the logic by which embryos decode morphogen inputs remains incompletely understood. Most models have focused on spatial gradients and thresholds, exemplified by Wolpert’s “French flag” model^2^, yet accumulating evidence suggests that timing and duration are equally critical. However, defining the exact cell fate requirements has been hampered by the lack of tools to precisely manipulate morphogens in both the spatial and temporal dimensions. Thus, a central open question is whether there exist minimal, experimentally testable ‘rules’ for when, where, and for how long morphogens must act to establish tissue architecture.”

During gastrulation, the pluripotent epiblast differentiates into the three germ layers (ectoderm, mesoderm, and endoderm), establishing the blueprint for embryonic development. Morphogen signaling, particularly through the BMP, WNT, and NODAL pathways, orchestrates this process^3,4^. BMP signaling from extraembryonic tissues initiates a signaling cascade that activates WNT and subsequently NODAL in the epiblast, collectively breaking anterior-posterior symmetry and specifying mesodermal fate (**Fig 1A**)^1,4,5^. While morphogen signaling pathways guide development, we still lack a precise understanding of how their spatiotemporal dynamics encode cell fate decisions and organize the germ layers.

**Fig 1.**
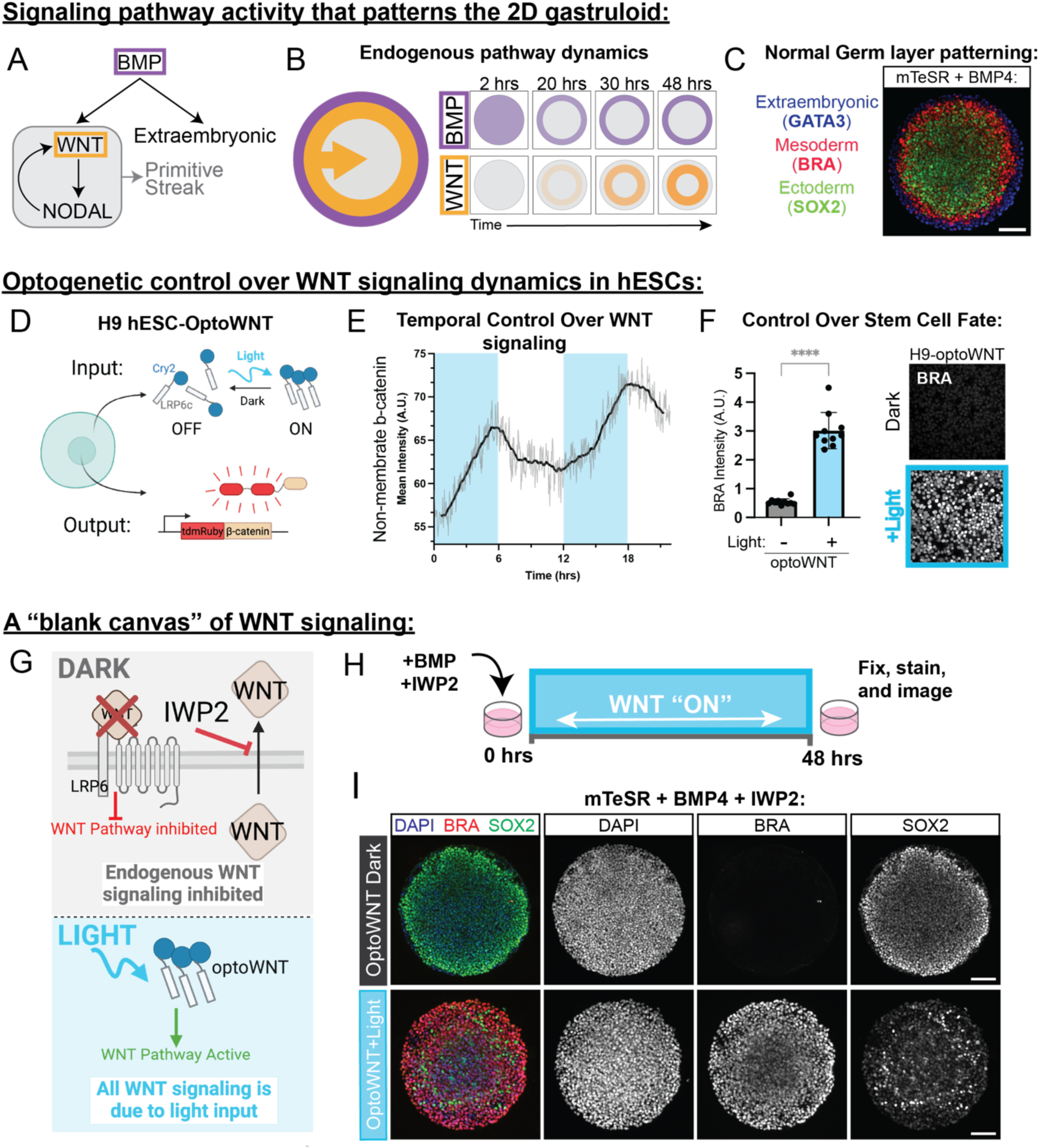
An optogenetic platform that controls WNT signaling dynamics in the 2D gastruloid. All scale bars are 100 µm. **(A)** Proposed hierarchy of signaling that patterns the 2D gastruloid. Exogenous BMP4 activates WNT signaling and activates NODAL in a feedforward loop that becomes self-sustaining. BMP signaling gives rise to the extraembryonic fates while WNT and NODAL signaling give rise to the primitive streak. **(B)** Schematic illustrating endogenous BMP4 (purple) and WNT (orange) signaling dynamics in the 2D gastruloid, that give rise to normal germ layer patterning. BMP4 signaling (SMAD) is restricted to the colony edge by 12 hours after addition of BMP, while WNT signaling (β-catenin) increases over time in a ring a few cell widths in from the colony edge, that slowly propagates inward over time. **(C)** H9-optoWNT hESCs plated in micropattern culture on 500 µm discs differentiated with BMP4 for 48 hours (without optogenetic activation or light exposure) and stained for fate markers, GATA3, BRA, and SOX2. Imaged at 20X. **(D)** Schematic of H9-optoWNT cell line expressing stably integrated opto-LRP6 to control WNT signaling in response to blue light, and CRISPR tagged β-catenin live cell reporter to visualize WNT signaling in hESCs. **(E)** Quantification of non-membrane β-catenin in response to light over time. DMDs were used to activate OptoWNT in 6-hour increments. Blue blocks represent applied light stimulus; no blue block indicates no optogenetic activation. **(F)** Antibody stain for Brachyury in H9-OptoWNT hESCs with and without light stimulation for 24 hours. Quantification 10 fields of view, imaged at 40X. P value calculated with an unpaired t test in prism (P<0.0001). **(G)** Schematic illustrating the “blank canvas” of WNT signaling created by the addition of the WNT ligand secretion inhibitor IWP2 in the 2D gastruloid. IWP2 inhibits all endogenous WNT signaling induced by the addition of BMP4, and WNT signaling is added back in with light, activating optoWNT downstream of the receptor. **(H)** Schematic representing the experimental design for optogenetic differentiation of blank canvas gastruloids shown in I (*bottom*). **(I)** Germ layer antibody stain for H9-optoWNT hESCs plated in micropattern culture and treated with BMP and IWP2 in the dark (*top*) and stimulated with blue light for 48 hours (*bottom*). Antibodies are Brachyury (red), SOX2 (green) and nuclear marker (blue).

Due to ethical and experimental limitations associated with studying human embryos, stem cell-derived embryo models have emerged as powerful tools to investigate developmental mechanisms^6–9^. Among these models, two-dimensional micropatterned human pluripotent stem cell colonies, termed “2D gastruloids,” stand out for their ability to recapitulate the cell fate decisions that occur during gastrulation in the human embryo as well as their exceptional reproducibility and experimental tractability^10–12^. As a result, 2D gastruloids provide a quantitative and high-throughput system for probing how spatiotemporal signaling dynamics shape cell fate decisions during early embryonic development. Stimulation of 2D gastruloids with exogenous BMP4 cause them to form concentric rings of germ layers and extraembryonic lineages with reproducibility sufficient to detect teratogenic effects through changes in spatial patterning^13^. Unlike heterogeneous 3D embryo-like aggregates^14–17,^ 2D gastruloids enable precise control over initial colony geometry and boundary conditions, enabling systematic exploration of the role of morphogen signaling dynamics on patterning.

Upon induction with exogenous BMP4 ligand, BMP signaling in the 2D gastruloid is first active throughout the entire colony, but due to receptor accessibility and inhibitor production is then restricted to the colony edge^18–20^. WNT signaling follows BMP activity and propagates as a wave (**Fig 1B**), moving from the edge to the center of the colony^5^. While multiple models of morphogen signaling^5,13,21,22^ have suggested the importance of the timing and duration of signaling waves, a systematic analysis of the role of morphogen dynamics in patterning that gastruloid has been technically challenging. However, these studies necessarily relied on endogenous signaling and global pharmacological perturbations, making it impossible to independently program WNT onset, duration, and spatial domain. As a result, the minimal spatiotemporal requirements sufficient to restore germ-layer organization have remained undefined.

Optogenetic control of developmental signaling provides this capability by enabling nearly instantaneous, reversible, and spatially precise activation of developmental signaling pathways. Recent advances in optogenetic tools, including OptoBMP^23^, OptoSOS (ERK signaling)^24,25^, and OptoWNT^26^, highlight their power for dissecting complex cell fate decisions by enabling the precise manipulation of signaling dynamics previously inaccessible with conventional ligand stimulation. Thus, optogenetics is ideally suited to systematically unravel how signaling dynamics shape embryonic development. Indeed, ‘optogenetic rescue’ experiments that define the minimal signaling requirements of Erk in morphogenesis successfully rescued an embryonic-lethal patterning mutant in *Drosophila*^27^. Although recent studies validated the utility of OptoWNT in directing mesoderm differentiation in human pluripotent stem cells^26^, how spatiotemporal regulation of WNT signaling shapes gastruloid patterning and fate decisions remains poorly understood. Here, we address this gap by developing an optogenetic “blank-canvas” approach that inhibits endogenous WNT secretion and reintroduces WNT pathway activity with light, allowing us to test sufficiency directly by programming onset, duration, and spatial location.

Here, we develop an optogenetic ‘blank-canvas’ gastruloid in which endogenous WNT secretion is blocked and pathway activity is reintroduced using a light-gated LRP6 receptor. Systematic temporal scans reveal a narrow WNT competence window during which onset and duration jointly control mesoderm output and tissue architecture, with the timing of this window set by cell density and enhanced by BMP priming. Using single-cell RNA sequencing, we show that WNT timing not only reorganizes germ layers but also redirects mesodermal subtype specification. Finally, by restricting WNT to a mid-ring within this window, we achieve a full optical rescue of germ-layer patterning with boundary fidelity that surpasses ligand-based induction. These findings establish WNT as a temporal morphogen and provide a general strategy to read out and engineer temporal signaling codes in embryo models.

## Results

### A high-throughput platform to deconstruct WNT pathway dynamics in the 2D gastruloid

To simultaneously visualize and control WNT signaling dynamics, we constructed an H9 hESC cell line (H9-optoWNT) bearing transposon integrated optogenetic LRP6, the WNT ligand coreceptor, and CRISPR-tagged tdmRuby3-β-catenin (**Fig 1D**). We confirmed that this edited cell line maintained PSC morphology and expressed pluripotency markers at comparable levels to unedited H9 hESC (**fig S1A,B**). When canonical Wnt signaling is active, β-cat accumulates in the nucleus, which results in the transcription of Wnt target genes, and in hESCs, differentiation into mesoderm as indicated by the mesoderm fate marker Brachyury (BRA). To ensure dynamic control over WNT signaling, we live imaged tdmRuby3-β-catenin while stimulating the optogenetic tool with blue light (405 nm) **Fig 1E**). We applied and removed the light stimulus in increments of 6 hours and quantified non-membrane β-cat in single cells in response to blue light stimulation (blue boxes indicate light stimulus). β-cat increased when the light stimulus was applied and decreased as soon as the stimulus was removed demonstrating the ability to reversibly control WNT signaling dynamics with light. To understand how optogenetic WNT activation compared to standard small molecule activation, we monitored β-cat translocation in OptoWNT cells stimulated with blue light and cells treated with CHIR 99021 for 6 hours. Blue light stimulation drove non-membrane β-cat to comparable levels to CHIR treatment (**Fig SG,H**). We then asked if opto-Wnt stimulation could drive hESCs into the mesoderm fate. We stimulated opto-Wnt cells and WT control H9 hESCs each seeded at 1,500 cells/mm^2^ with blue light (405 nm) for 24 hours and fixed and stained for the mesoderm fate marker BRA. Compared to the control, H9-optoWNT cells differentiated into the mesoderm fate as indicated by the expression of the BRA fate marker (**Fig 1F, fig S1C**). Notably, there was no detectable dark state activation in the optogenetic cell line. Additionally, we confirmed the ability of H9-optoWNT to direct transcription of WNT target genes in response to light (**fig S1D**). Overall, these experiments demonstrate that H9-OptoWNT mimics canonical WNT signaling in hESCs and directs WNT driven differentiation.

Micropatterned stem cells treated with BMP4 for 48 hours form concentric rings of germ layers (endoderm, mesoderm and ectoderm) as well as an outer extraembryonic amnion-like layer (**Fig 1C, S2A**)^10^. We selected 500 µm micropatterns because they reliably generate these germ layers while allowing for multiple replicates per well (>40 gastruloids per well of a 96-well CYTOO plate). Since WNT ligands are produced during the differentiation due to the addition of exogenous BMP4,^28^ we sought an approach that would allow precise control over the amount of WNT signaling in the system by blocking endogenous WNT signaling. We introduced IWP2, a WNT secretion inhibitor^29,30^ (**Fig 1G**), effectively removing endogenous WNT signaling and creating a “blank canvas” for controlled optogenetic activation. Consistent with WNT’s essential role, gastruloids co-treated with BMP4 and IWP2 do not form the germ layers (**Fig 1I**, *top*). Optogenetic activation of WNT with constant illumination rescued mesoderm differentiation (quantification of mesoderm rescue in **fig S2B**) but disrupted normal tissue patterning, expanding the mesoderm pattern across the entire colony, eliminating the outer GATA3+ extraembryonic-like ring (**fig S2C**) and significantly reducing the size of the SOX2+ ectoderm inner ring (**Fig 1I**, *bottom*). These results indicate that continuous, uniform WNT activation is insufficient to rescue proper gastruloid patterning, highlighting the need for precise temporal regulation. To systematically define the temporal requirements for WNT-driven mesoderm induction and tissue organization, we varied the duration and onset of global WNT activation using optogenetics.

### Optogenetic screen reveals temporal requirements for mesoderm patterning in 2D gastruloid

#### Both the onset and duration of WNT signaling control mesoderm differentiation

Having established that uniform and constant activation of WNT in our “blank canvas” gastruloid system does not rescue patterning in the gastruloid, we next sought to determine how the timing and duration of WNT signaling influence mesoderm specification. While live cell timelapse microscopy of WNT signaling during germ layer formation in the 2D gastruloid reveals that β-catenin increases in a ring that slowly propagates toward the colony center,^5^ it remains unclear how temporal features of WNT activity contribute to cell fate outcomes. To systematically investigate this, we used global optogenetic illumination with defined durations and onset delays to map the temporal requirements for WNT-driven mesoderm induction and pattern organization.

Using our high-throughput LITOS (LED Illumination Tool for Optogenetic Stimulation) device^31^, we systematically explored how both the duration and timing of WNT signaling affect mesoderm patterning in the 2D gastruloid. We varied WNT from 6 to 48-hours and tested initiation delays from 0 to 42 hours (**Fig 2A**, *schematic*). To ensure that WNT signaling came solely from our optogenetic activation, we added IWP2 to inhibit endogenous ligand production. After 48 hours, we stained for germ-layer markers and averaged the colony morphology across 10 replicates per condition (**Fig 2A**). The DAPI channel is consistent across all treatment conditions (**FigS3A**).

**Fig 2.**
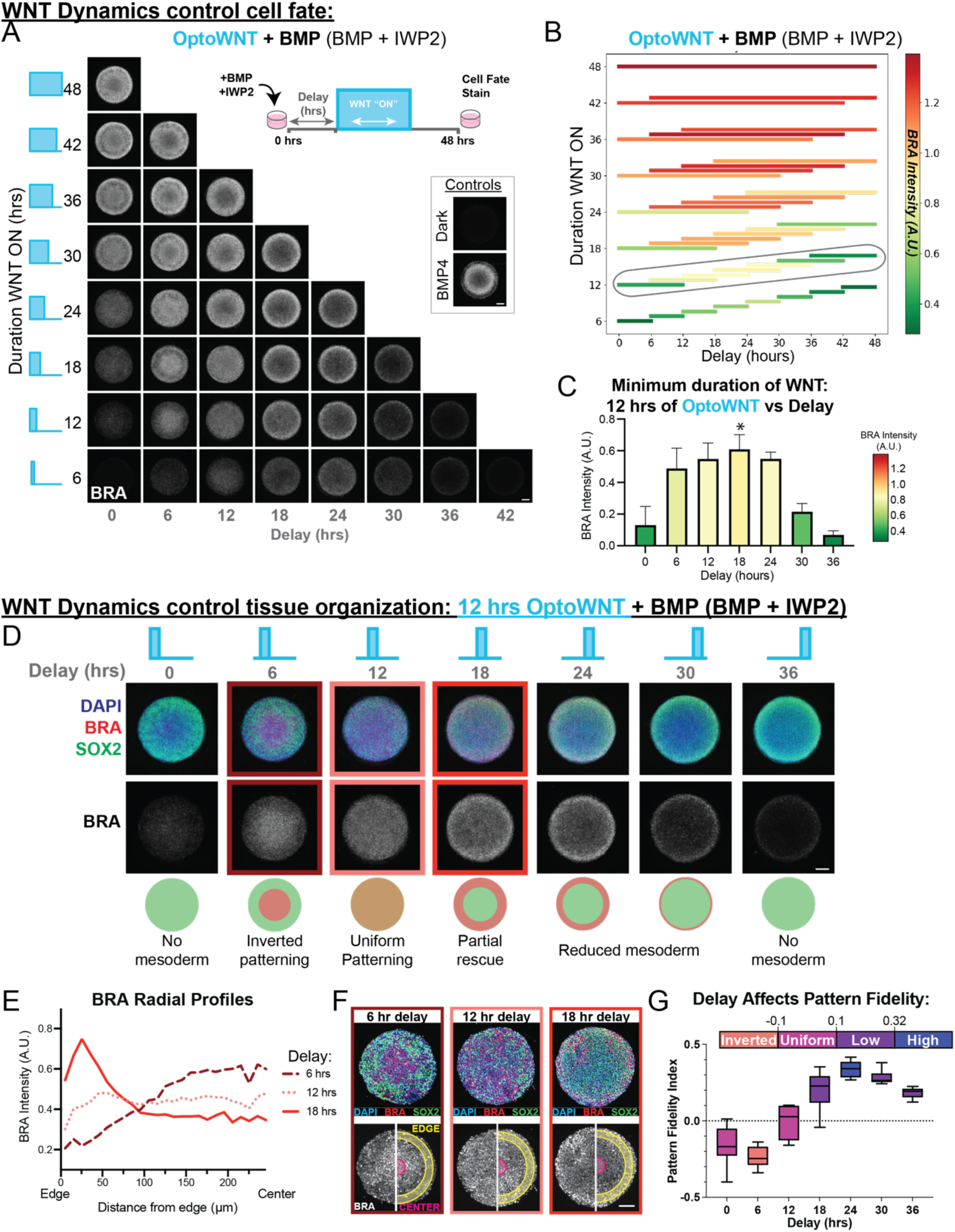
WNT timing and duration modulate mesoderm differentiation and tissue organization in the 2D gastruloid. Immunofluorescence results from an optogenetic screen to identify the minimum duration of WNT required to pattern the gastruloid using the LITOS high throughput LED light delivery device. LITOS stimulation conditions: 1s on every 20s. Micropatterns were treated with BMP and IWP2 to create a “blank canvas” of WNT signaling so all WNT activation is solely due to activation of the opto tool. A BMP4 treated well and a BMP4+IWP2 dark treated well were included as controls. All scale bars are 100 µm. **(A)** Both the duration (length of time WNT was activated) and the delay (length of time prior to WNT activation) were varied (*schematic*) from 0 to 48 hours in 6-hour increments. Treatments were added at T0 and samples were fixed and stained for fate markers at 48 hours. The colony average of 10 representative colonies stained for Brachyury is shown for each light treatment condition, and for each control. **(B)** Heat map bar plot representing the average mesoderm differentiation for each light treatment over 10 replicate colonies shown in 2A. The color of each bar represents the intensity of Brachyury (red is the highest intensity and green is the lowest). The length and location of each bar represents the duration and timing of the WNT signal respectively. The mean level of mesoderm differentiation per light treatment was quantified based on DAPI nuclear segmentation. **(C)** BRA intensity for gastruloids treated with 12 hrs of OptoWNT. IF data shown in 2A. Asterisk (*) denotes the “partial rescue” conditions: 12 hrs of OptoWNT with an 18-hour delay. This is the delay prior to WNT activation that achieves the highest amount of mesoderm differentiation for the minimum duration. D through G: all data from gastruloids treated with 12 hours of OptoWNT. **(D)** Merged immunofluorescence colony averages (10 replicate colonies) antibody stained for SOX2 (green), Brachyury (red), and nuclear marker (blue) for 12 hours of WNT signaling at each delay demonstrating differences in tissue organization depending on timing of WNT. **(E)** Radial profiles (Brachyury intensity as a function of distance from the edge) for 12 hours of WNT with a 6-, 12-, and 18- hour delays. **(F)** Representative colonies for treatments with radial profiles shown in (E) demonstrating differences in patterning. Pattern fidelity index is calculated using the edge and center regions (*bottom*). **(G)** Pattern fidelity index calculated using the average BRA intensity in the 11-40 µm region as the edge, and the 211-240µm region as the center (distance from edge). Box plot is colored according to the degree of pattern complexity (high, low, uniform and inverted patterning).

To summarize these data, we quantified the average mesoderm intensities over 10 replicate colonies and created a visualization where each bar represents the duration of WNT signaling (y axis), when the WNT signal was received (x axis), and resulting mesoderm intensity (color-coded, red indicating highest intensity) (**Fig 2B**). While mesoderm differentiation generally increased with longer durations of WNT (also in **fig S3B**), notably, identical durations produced vastly different differentiation levels depending on when signaling began (**Fig 2B**). For example, 6 hours of WNT delivered with a 24-hour delay results quantifiable mesoderm differentiation while 6 hours of WNT delivered at 42 doesn’t result in any detectable mesoderm differentiation (**FigS2D**). These findings demonstrate that both the timing and duration of WNT activity determines the amount of mesoderm differentiation.

#### The minimum duration of OptoWNT required to rescue mesoderm differentiation is 12 hours with an 18-hour delay

To identify the minimum duration of WNT signaling required to restore the same level of mesoderm differentiation induced by BMP4 alone, we normalized the data to the average mesoderm intensity for the BMP4 control colonies and found that 12 hours of WNT delivered with an 18-hour delay achieves the target amount of mesoderm differentiation for the minimum WNT input (**Fig 2C**). We refer to these minimum temporal requirements as the “partial rescue” for mesoderm differentiation.

Next, we examined germ layer organization of the optically patterned colonies for light treatment conditions that received 12 hours of WNT by visualizing both SOX2 (green) and BRA (red). We observed striking differences in tissue organization depending on when WNT signaling began. For instance, a 6-hour delay resulted in inverted patterning, with SOX2+ ectoderm surrounding central BRA+ mesoderm. A 12-hour delay disrupted typical concentric ring formation entirely, producing uniformly distributed SOX2+ and BRA+ cells across the colony (**Fig 2D**).

The organizational differences are clearly captured in the radial intensity profile (**Fig 2E**). In control colonies treated with BMP4 alone, brachyury intensity appears just inside the extraembryonic outer ring (between 50-100 µm from the colony edge) and decreases in the center of the colony (**fig S3C**). This profile also appears in the 18 hr delay “partial rescue” condition, though the BRA peak shifts toward the colony edge (0-50 µm) (**Fig 2E**, *solid line*). Conversely, colonies with a 6-hour delay between BMP4 + IWP2 addition and OptoWNT activation showed an inversion of the profile, with the region of max intensity in the center of the colony (**Fig 2E**, *dashed line*). Whereas, a 12-hour delay abolished the radial symmetry profile entirely, resulting in a uniformly flat intensity profile across the entire colony (**Fig 2E**, *dotted line*).

To quantitatively capture these differences in radial organization we developed the Pattern Fidelity Index (PFI). The PFI quantifies how well BRA expression is localized to its expected radial domain by comparing the BRA intensity at the expected max near the colony edge (**Fig 2F**, *yellow ring*) and the expected minimum at the colony center (**Fig 2F**, *magenta*). Notably, BMP4 patterned colonies achieve a PFI value of greater than 0.32 while low values (0.1–0.32) indicate less radial symmetry and negative values (less than -0.1) indicate inverted patterning. Further details on PFI calculation and interpretation are provided in Supplemental Figure 4 and the Methods section (Pattern Fidelity Index Metric).

We quantified the PFI for each delay of the 12 hours of OptoWNT treated colonies. Remarkably we observed that the full spectrum of tissue organization outcomes can be achieved from the same duration of WNT: inverted patterning (6 hr delay), uniform patterning (12 hr delay), low pattern fidelity (delays 18, 30 and 36 hrs), and even high pattern fidelity (24 hr delay) (**Fig 2G**). These finding demonstrate that the timing of WNT, plays a role in determining how the tissues are organized. Overall, these results highlight that WNT signaling during the middle timepoints of gastruloid differentiation (roughly between 12 and 24 hours after BMP addition) is essential for both the total amount of mesoderm differentiation and the correct spatial organization of germ layers. Recently, intermediate durations of BMP signaling were shown to optimally induce mesoderm fate in unpatterned hESCs^32^. Therefore, we asked whether the duration of BMP treatment prior to WNT activation explained the window of maximal mesoderm differentiation here.

### BMP and cell density regulate the critical period for WNT induced mesoderm differentiation

#### Mesoderm differentiation is controlled by the timing of WNT signaling without BMP

To test how the gastruloid is patterned by activation of WNT alone, we repeated the optogenetic delay-duration screen without BMP4. Briefly, we illuminated IWP2-treated micropatterns with durations of OptoWNT activation from 6 to 48 hours and tested the same delays explored above (**Fig 3A**, *schematic*). If the delay between BMP signaling and WNT activation is responsible for the critical window of mesoderm differentiation, then WNT duration should produce the same amount of mesoderm regardless of its timing. On the other hand, if a colony-intrinsic factor or parameter that changes as a function of time after plating, such as cell density, also contributes to the critical window then the critical window should still exist. We observed the same critical window even in the absence of BMP: WNT activity delivered at intermediate timepoints resulted in maximal mesoderm (**Fig 3A**, **fig S5**). These results suggest that the timing of the critical window is set by a colony intrinsic factor.

**Fig 3.**
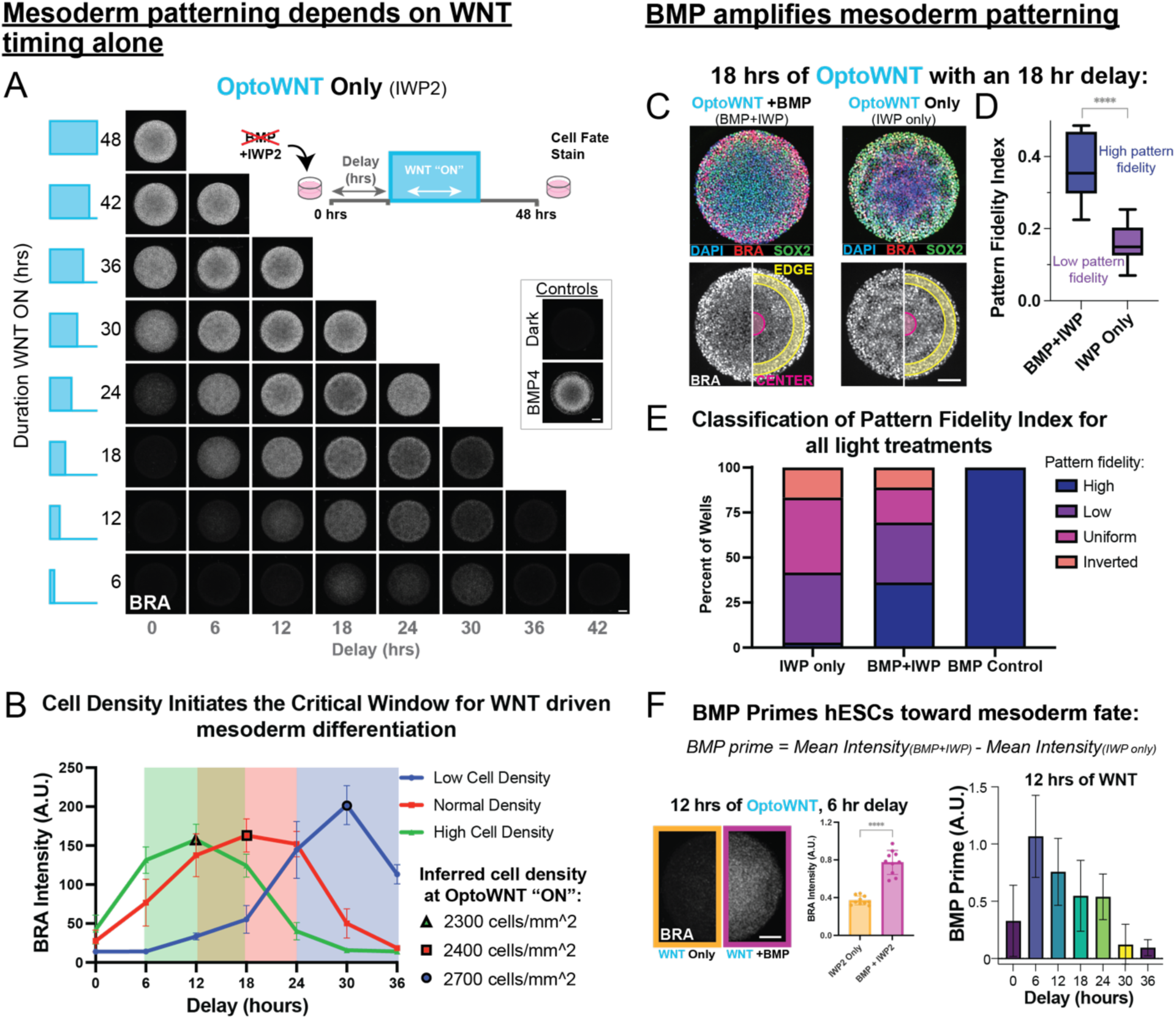
BMP and cell density regulate the critical period for WNT induced mesoderm differentiation. **(A)** Immunofluorescence results from the same experiment in figure 2, but without BMP4 (micropatterns treated with IWP2 only). IF Colony average of 10 representative colonies stained for Brachyury for each light treatment condition, and controls. **(B)** Quantification of immunofluorescence imaging of un-patterned (monolayer) OptoWNT hESCs differentiated with 12 hours of WNT with the depicted delay. Cells were plated at different densities to test the critical window for mesoderm differentiation. Imaging was done at 40X and 20 fields of view were collected and factored into quantification of each point in the plot. Cells were segmented on DAPI channel. Highlighted points (black outline): inferred cell density when optoWNT was activated, resulting in maximal mesoderm differentiation. **(C)** Representative IF colonies for micropatterns differentiated with 18 hours of WNT delivered with an 18-hour delay and treated with either BMP and IWP2, or IWP2 alone. **(D)** Pattern fidelity index calculated for 10 representative colonies with and without BMP. **** denotes P<0.0001. **(E)** PFI Classification for all light treatments from figures 2A and 3A, treated with either IWP2 alone, or BMP and IWP2, compared to BMP4 control wells. **(F)** *Far left*: Example mean colony images of treated with the same duration and delay of OptoWNT, with and without BMP, demonstrating the BMP priming effect. *Center:* average mesoderm quantification for the 10 colonies shown (*left*). Unpaired t test P< 0.0001. *Right*: BMP prime quantified by the difference between the mean Brachyury intensity for colonies treated with BMP and IWP2, and IWP2 only for gastruloids treated with 12 hours of OptoWNT at each delay.

#### Cell Density initiates the critical period for WNT induced mesoderm differentiation

We tested if this effect was micropattern-independent by altering the delays on unpatterned hESCs and found the same window was maintained (**fig S5E**) demonstrating that the critical window is not due artificial confinement on the micropatterned surface. To determine if this critical window was a function of cell density, we seeded cells with different initial cell densities representing the range in density of a typical 48-hour experiment. We found that shifting the initial seeding density is sufficient to alter the time of peak mesoderm differentiation (**Fig 3B**). A lower initial cell density moved the critical window to later delays (**Fig 3B**, *blue*), and a higher cell density shifted the window to earlier delays (*green*) compared to the control initial seeding density (*red*).

To determine the optimal density for mesoderm differentiation, we estimated cell density over time during the 48 hour differentiation (**Supp Table 2**) based on published H9 hESC doubling times^33,34^ (**Fig 3B**). The cell density at the onset of OptoWNT activation that achieved the maximal mesoderm differentiation was maintained regardless of starting density revealing that WNT signals delivered when the cells were at an inferred density of 2400 cells/mm^2^ resulted in peak mesoderm differentiation.

Overall, this implicates cell density as a crucial regulator for the critical window of mesoderm differentiation. This result also suggests that the reason the intermediate durations of BMP achieved maximal mesoderm differentiation reported by Camacho-Aguilar et al., may be because cell density was optimal for WNT signal interpretation into mesoderm in that window^32^. Indeed, WNT signaling-induced mesoderm differentiation has been shown to depend on cell density in many contexts^20,32,35^, and our work demonstrates that this dependence results in a critical window for optimal mesoderm fate determination in hESCs and the 2D gastruloid. This is consistent with literature demonstrating that both ligand-^32^ and CHIR-^36^ induced nuclear β-cat translocation is sensitive to cell density.

#### BMP increases both gastruloid pattern fidelity and total amount of mesoderm

While BMP4 was not responsible for the critical window for mesoderm differentiation, this experiment revealed a different role for BMP4 in patterning OptoWNT-treated gastruloids. Mesoderm patterning was much closer to a normally patterned 2D gastruloid for BMP + OptoWNT treated colonies than without BMP. For example, 18 hrs of OptoWNT delivered with an 18-hour delay without BMP did not produce the expected outer ring of max BRA intensity, but rather BRA signal was uniform across the entire colony (**Fig 3C**, *right*). However, when BMP was present, the expected radial geometry was recovered, with an outer mesoderm band and inner SOX2+ ring (**Fig 3C**, *left*). This was reflected in the PFI, with BMP + OptoWNT achieving high pattern fidelity while OptoWNT alone results in low pattern fidelity, despite receiving the same duration and delay of WNT (**Fig 3D**).

To explore the role of BMP4 in pattern fidelity more generally, we calculated the PFI for all treatments included in both duration-delay screens (+BMP & -BMP). We found that when treated with BMP, OptoWNT differentiated gastruloids had a PFI comparable to control colonies in greater than 35% of light treatments, while OptoWNT alone never achieved a PFI equivalent to control gastruloids (**Fig 3E**). Interestingly, treating micropatterns with WNT3A ligand alone has been shown to be sufficient to induce the mesoderm band with high fidelity in the 2D gastruloid^1^. Therefore, our results suggest that the recovery of a mesoderm band upon WNT3A application does not deliver truly uniform WNT pathway stimulation and in fact induces non-uniform activation or inhibitor production. Thus we can conclude that truly uniform activation of WNT is not sufficient to pattern the gastruloid^28^.

In addition to increasing the pattern fidelity we noted that BMP4 also increases the total BRA produced in the colony as a function of WNT. Comparing optogenetically patterned gastruloids differentiated with OptoWNT alone to those differentiated BMP+OptoWNT revealed that BMP4 increases the total amount of mesoderm differentiation (**Fig 3F, fig S6**). For example, comparing all delays/timepoints tested for 12 hours of WNT, there is a statistically significant increase in BRA intensity (P=0.0095) when patterns are treated with BMP4 and IWP2 compared to IWP2 alone (**fig S6B**). Thus, BMP4 increases the total amount of mesoderm differentiation through some other mechanism besides WNT ligand secretion. TGFβ ligands have been reported to enhance the response of β-catenin through a mechanism independent of WNT ligand induction.^37^ However the downstream effect on stem cell differentiation and germ layer patterning has not yet been reported. Therefore, it is possible that BMP enhances the nuclear translocation of β-catenin to an extent sufficient to influence the level of mesoderm differentiation observed here.

Overall, these experiments demonstrate that BMP4 plays a role in enhancing both the pattern fidelity and total amount of mesoderm differentiation in the 2D gastruloid. While BMP4 does not define the critical temporal window for mesoderm induction, it substantially improves radial patterning accuracy when combined with WNT activation. These findings highlight a previously unexplored role for BMP4 in amplifying mesoderm differentiation. Having identified these essential signaling requirements, we next sought to determine whether a correctly timed optogenetic WNT stimulation could faithfully recapitulate the set of cell types produced by standard BMP-induced differentiation.

### Optogenetic WNT activation with optimal timing produces the same cell types as BMP induced differentiation

After identifying the minimal conditions required for partial optical rescue of mesoderm patterning, two key questions remained unresolved: first, whether the cell identities generated in optogenetically patterned colonies fully recapitulate those produced by standard BMP4 induction or merely express similar fate markers; and second, whether aberrantly patterned colonies represent normal cell identities organized differently, or instead comprise entirely distinct cell types. To address these questions and further elucidate the mechanisms underlying fate specification in the 2D gastruloid, we performed single-cell RNA sequencing.

We chose four optogenetically-induced Wnt delays (0, 6, 12, and 18 hours). These included the 12 hours as the duration required to rescue mesoderm patterning **(Fig 2C**) and conditions that result in aberrant patterning phenotypes (0 hr delay: no mesoderm, 6 hr delay: inverted patterning, and 12 hr delay: uniform patterning) (**Fig 4A**). Gastruloids treated with BMP4 alone served as a positive control, and BMP4 + IWP2 in the dark served as our negative control. Each condition was analyzed using single-cell RNA sequencing alongside corresponding immunofluorescence to validate cell identity and pattern fidelity.

**Fig 4.**
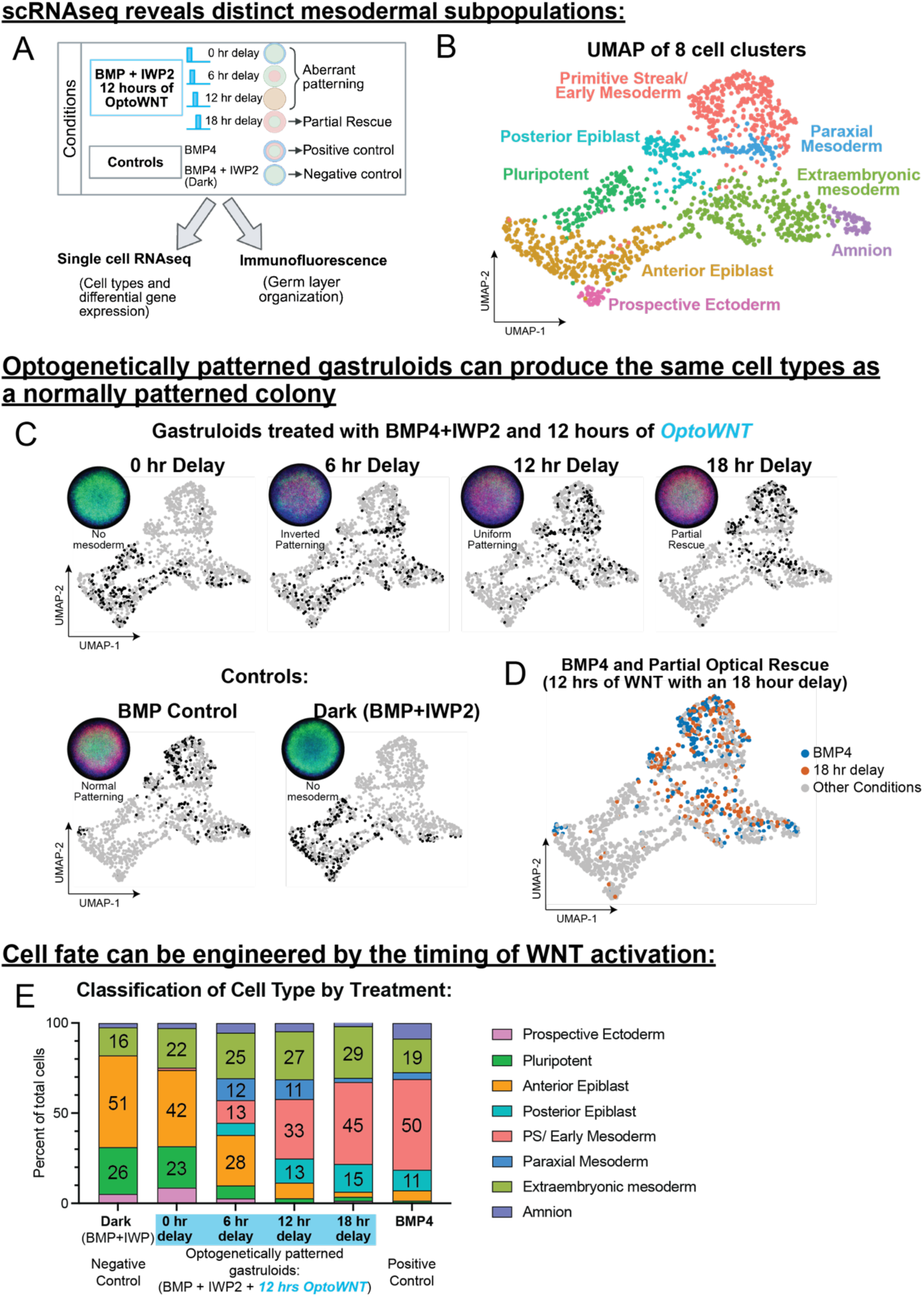
Optogenetic WNT activation with optimal timing produces the same cell types as BMP induced differentiation. **(A)** Schematic demonstrating the scRNAseq experiment workflow. Four light treatment conditions that all received the same total duration of WNT (12 hrs) were chosen for this analysis; three revealed aberrant patterning phenotypes in the high throughput optogenetic screen (0-, 6-, and 12-hour delays). The fourth light treatment condition was the optimal condition for the minimum duration of WNT required to rescue mesoderm differentiation (18-hour delay). All optoWNT conditions were treated with BMP and IWP2 to create the blank canvas of WNT signaling. Undifferentiated (dark) BMP+IWP2 and BMP4 treated gastruloids were included as controls. H9 optoWNT hESCs were plated in duplicate in micropattern culture and treated with the 4 different drug and light treatment combinations. After 48 hours, one well of each treatment was lifted for scRNAseq, and one well was stained for germ layer markers for corresponding germ layer organization analysis. **(B)** UMAP projection of 8 cell clusters resulting from the integrated dataset (cells from all treatments) detected by unsupervised clustering in 2D gastruloids. **(C)** Highlighted cells from each treatment in the UMAP space, and corresponding immunofluorescence average for that treatment. IF image is an average of the merged image (DAPI-blue, BRA-red, SOX2-green) of 10 replicate colonies from that treatment. *Top*: light treatment wells. *Bottom*: control wells. Note: Since SOX2 expression is so high in 0 hr delay, and Dark control, the SOX2 LUTS are shown at a lower setting than the other treatments so patterning phenotypes can still be visualized. Merged colonies and individual channels all set to the same LUTS are shown in supplemental fig7A. **(D)** Overlay of cells from the BMP4 control treated well and the partial rescue treatment (BMP+IWP2, with 12 hours of WNT delivered with an 18-hour delay) highlighted in Blue and Orange respectively. **(E)** Stacked bar plot illustrating the distribution of cell types for each treatment as a percent of total cells from that treatment.

After optimization, unsupervised clustering revealed 8 distinct clusters (**Fig 4B**). Using canonical fate markers, the top differentially expressed genes in each population, and publicly available gene expression data of gastrulating human embryos^38–40^, we identified these clusters as primitive streak/early mesoderm, a more advanced mesoderm population likely corresponding to paraxial and lateral plate mesoderm, two epiblast populations (one primed toward posterior fates and one primed toward anterior fates), a prospective neural ectoderm population, and two extraembryonic lineages (amnion-like and extraembryonic mesoderm-like cells) (**See fig S7C-E and supplemental Note 1 for explanation of cell type annotation**).

The cells from OptoWNT-treated gastruloids with a 0 hr delay fall in essentially the same UMAP space as the Dark control (**Fig 4C**). As the delay increases toward the partial rescue treatment (18 hr delay), the distribution of single-cells more closely resemble the BMP4 control (**Fig 4C**). Remarkably, the cells from the partial optogenetic rescue and BMP4 patterned colonies have nearly identical distribution in UMAP space (**Fig 4D**).

To better understand how each delay alters cell-type distribution, we quantified the proportions of the cell types as a percent of total cells. All treatments result in similar amounts of the extraembryonic lineages (**Fig 4E**) including the dark BMP+IWP2 condition, further supporting the extraembryonic identity of these clusters likely both arising from BMP4 signaling as opposed to cell types that require WNT signaling. Remarkably, the cell types from the Dark negative control and illuminated OptoWNT with a 0-hour delay result in nearly identical distributions, reinforcing the importance of the critical window. As the delay increases toward the partial rescue, the cell type distribution gets closer to BMP4 patterned colonies, with an 18-hour delay resulting in essentially the same cell types as the positive control.

Interestingly, gastruloids subjected to OptoWNT stimulation at 0- and 6-hour delays contained a substantial fraction of pluripotent cells and anterior primed epiblast, populations that are absent in both the BMP-patterned controls and the partial rescue (18-hour delay) condition yet present in the Dark control (**Fig. 4E**). Comparing the partial rescue condition to the BMP4 control demonstrates almost the exact same distribution in cell types, however, there is a slight reduction in amnion cells. Finally, when comparing the cell types between gastruloids that have aberrant patterning (6- and 12-hour delays) we noticed that a unique mesodermal population, paraxial mesoderm, emerges (**Fig 4E**, **fig S8**). Therefore, the cell types in aberrant patterning phenotypes (0-, 6- and 12-hour delays) are not only in a different organization as indicated by immunofluorescence analysis, but they also produce different BRA+ and SOX2+ cell types. These results underscore that disruptions in WNT signaling timing not only profoundly affect tissue organization but also shift cell identities toward alternative developmental trajectories.

### Full optical rescue of germ layer patterning requires temporal and spatial control over WNT signaling and exogenous BMP4

While the 18-hour delay rescued most of the cell types of the gastruloid, scRNA-seq revealed that very few extraembryonic Amnion-like cells were recovered in comparison to the BMP4 control (**Fig 4E**). Corresponding IF imaging of germ layer organization revealed that the GATA3-positive outer ring is lost in full-colony-illuminated OptoWNT cells (**Fig 5A**). In addition, the BRA ring of the partial rescue peaks slightly outside the peak of the BMP4 patterned colony, as evidenced by the lower PFI when quantified using the mesoderm ring location in a BMP4 patterned colony (**Fig 5B**).

**Fig 5.**
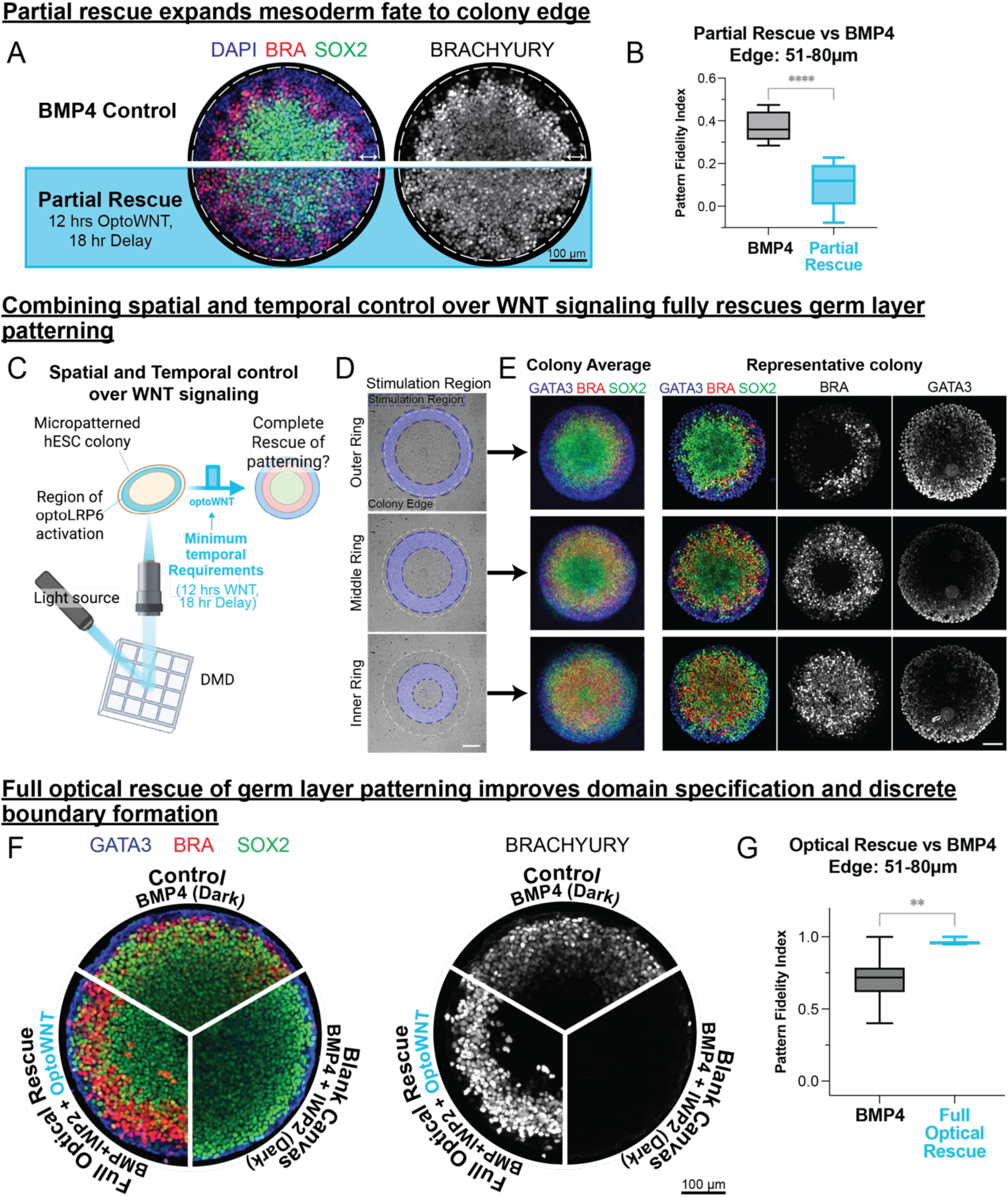
Application of optimal position, timing and duration of WNT results in full optical rescue of germ layer patterning. **(A)** representative colony for germ layer antibody stain (DAPI, BRA and SOX2) for gastruloids treated with BMP4 (*top*) and with BMP+IWP2 and the partial rescue conditions (blue box, *bottom*). White arrows show expected region of extraembryonic cells (GATA3+) that is substantially reduced in the optogenetic partial rescue gastruloids. **(B)** Comparison of the Pattern complexity index for the partial optical rescue (BMP+IWP2, 12 hours of WNT with an 18-hour delay) and BMP4 patterned colonies calculated from the target mesoderm edge region. ****P<0.0001 **(C)** Proposed experimental approach for a full optogenetic rescue of germ layer patterning in the 2D gastruloid. Using the DMDs engineered into the light path of our microscope, we control the location of OptoWNT activation in the colony, for the previously identified minimum temporal requirements (12 hours of WNT with an 18-hour delay). **(D)** DMD mask of the three possible optical rescue stimulation regions overlay with brightfield image of micropatterned hESCs. Dotted lines highlight the location of the stimulation region (blue) in relation to the edge of the colony (white). **(E)** Immunofluorescence results from stimulation with the corresponding stimulation region in E. Antibody stained for Brachyury (mesoderm, red), SOX2 (ectoderm, green) and GATA3 (extraembryonic, blue). *Left*: colony average of 7 replicate colonies and *Right*: representative colony, treated with the indicated stimulation region. **(F)** Comparison of location of germ layer (IF) stain for gastruloids treated with the indicated treatments. H9-optoWNT gastruloids treated with BMP4 (*top*) form discrete regions of GATA3 (ExE), BRA (mesoderm) and SOX2. Mesoderm is eliminated in H9-optoWNT gastruloids treated with BMP4 and IWP2 (*bottom right*). H9-optoWNT gastruloids treated with BMP4 and IWP2 and the minimum spatiotemporal requirements form discrete regions of GATA3 (ExE), BRA (mesoderm) and SOX2 comparable to BMP4 induced patterning. **(G)** Pattern complexity index quantified for BMP4 control patterned colonies compared to the colonies resulting from the optical rescue (8 colonies quantified from each treatment). See methods for quantification approach. **P=0.0037

In our high-throughput WNT duration experiments, the illumination spanned the entire area of each micropatterned colony, leading to colony-wide WNT activation. Although this global activation successfully restored the target mesoderm differentiation level (**Fig 2C**) and produced the full complement of gastruloid cell types (**Fig 4E**), it compromised the formation of distinct germ-layer boundaries and resulted in the loss of a clearly defined extraembryonic ring. We therefore hypothesized that precise spatial control of WNT signaling would enable full rescue of germ layer organization.

To test this hypothesis, we used digital micromirror devices (DMDs) integrated into our microscope’s optical path to spatially control activation of WNT signaling within gastruloids (**Fig 5C**). We evaluated three candidate illumination patterns spanning the area of the gastruloid (**Fig 5D, fig S9D**). Stimulation of the outer ring led to diminished mesoderm formation and incomplete radial symmetry yet interestingly preserved extraembryonic-like differentiation (**Fig. 5E**, *top*). Conversely, inner-ring illumination disrupted patterning, producing aberrant localization of SOX2+ cells between the extraembryonic and mesodermal domains, along with a central, undersized mesoderm region (**Fig. 5E**, *bottom*). Strikingly, illumination of the “goldilocks” region (about 32 µm from the edge), combined with the minimal temporal requirements identified previously (12 hours of WNT activation following an 18-hour delay), fully restored proper germ-layer organization, producing an outer extraembryonic ring of appropriate dimensions along with distinct and correctly positioned mesoderm and ectoderm bands (**Fig. 5D,E** *middle*). Notably, stimulating in this ring with nonoptimal temporal requirements (36 hrs of OptoWNT, 12 hr delay) does not fully rescue patterning, demonstrating that spatial control of WNT alone is not sufficient to fully rescue germ layer patterning, and both spatial and temporal control are required (**fig S9D**). We conclude that spatially restricted WNT activation in the middle ring is essential for accurate mesoderm differentiation and proper gastruloid patterning.

Finally, we computed the PFI of micropatterns treated with spatiotemporally patterned WNT activity (**Fig 5G**). Compared to the BMP4 control, colonies illuminated in this goldilocks region had a PFI that was significantly higher than BMP4 treated colonies, demonstrating improved domain specification and boundary formation using optogenetics compared to traditional ligand-based patterning methods. These results demonstrate that precise spatial and temporal control of WNT signaling, in combination with exogenous BMP4, not only recapitulates but surpasses the fidelity of traditional ligand-based gastruloid patterning methods. Thus, our optogenetic approach provides a powerful and refined strategy for dissecting and reconstructing developmental signaling dynamics.

## Discussion

### Minimal temporal rules for Wnt-mediated germ layer patterning

Developmental signals are increasingly recognized to act not just as static gradients, but as dynamic patterns whose timing and history govern cell fate^24,41–44^. By optogenetically controlling WNT in a human gastruloid, we directly map the temporal ‘rules’ that link WNT exposure history to germ-layer architecture. Our experiments reveal that human pluripotent stem cells and thus, likely the human epiblast, follow strict temporal requirements for WNT signaling to induce proper germ layer patterning. Specifically, we identified a narrow critical window during which WNT activation robustly drives mesoderm differentiation in the 2D gastruloid. Both the timing (onset) and duration of WNT signaling determine the final pattern. If WNT is applied too early or too late, too brief or too long, gastruloids fail to pattern their mesoderm correctly. These findings underscore the concept of a “temporal morphogen,” wherein the temporal profile of a signal (e.g. when and how long it is present) can be as important as its concentration. Consistent with this idea, prior studies have shown that the duration of WNT signaling (and it’s downstream NODAL wave) is a key determinant of mesoderm induction^5^. Our results now define the minimal temporal rules for WNT-mediated germ layer patterning: there exists a discrete window of competency during which WNT signaling proportionally directs mesoderm differentiation, whereas WNT outside of this window or below the duration threshold is ineffectual in establishing the correct mesoderm pattern.

### Timing between BMP and WNT determines cell fate

An important aspect of these temporal rules is the relative timing of BMP and WNT signals, which emerged as a mechanism for encoding the mesoderm cell fate decision. We found that BMP treatment acts as a priming cue that dramatically enhances Wnt-induced mesoderm differentiation as well as pattern fidelity. This suggests that BMP signaling prepares cells to interpret subsequent WNT signals to promote mesoderm differentiation. Mechanistically, BMP priming likely elevates the responsiveness of pluripotent cells by inducing early mesendoderm genes or autocrine factors (including WNT itself) that synergize with the incoming WNT cue^45^. Intriguingly, despite uniform optogenetic WNT activation in the colonies, the presence of BMP4 positions the germ layers in the expected concentric rings for most light treatments (Fig 2A and Fig 3C-E). Therefore, the BMP priming may be one mechanism by which BMP provides positional information for mesoderm differentiation that doesn’t require a paracrine signal. This results in spatial information because the time between the initial BMP signal and the onset of the WNT wave is interpreted into different fates.

In our blank canvas gastruloids, changing the delay between BMP and WNT produced different cell fate outcomes. For example, WNT activation closer to BMP (6- and 12- hour delays) yielded distinct mesoderm subtypes by scRNAseq compared to a longer delay. Optimal WNT timing (18 hours following BMP4) preferentially induced gene programs associated with the “organizer” and early mesoderm (e.g., early primitive streak markers), whereas an early WNT pulse (6-12 hours after BMP4) biased cells toward more paraxial mesodermal or more advanced mesodermal fates^46,47^. Thus, beyond simply determining whether mesoderm forms, the relative timing of BMP and WNT serves as a code that tunes which type of mesoderm forms. This temporal coordination could mirror how the embryo sequentially generates different mesoderm lineages as the primitive streak matures. It also demonstrates that cells not only require the right signals but must receive them at exactly the right time to unlock the full spectrum of posterior fates. Timing between signaling pathway activation coordinates developmental processes such as somitogenesis^48^. Therefore, this study also advances an emerging picture of the importance of the relative timing *between* signals.

An important consideration here is the role of secreted inhibitors in gastruloid patterning. The secreted WNT inhibitor DKK was shown to play an essential role in positioning the primitive streak in the 2D gastruloid^28^. However, since DKK inhibits WNT signaling through binding to LRP5/6 on the cell surface^49^, and activation of WNT here occurs downstream of ligand binding, expression of DKK would not have an effect. Additionally, the DKK binding domain on LRP6 is in the extracellular region and we only express the cytoplasmic domain, so there is no way for DKK to bind to the optotool.

### Influence of cell density on signaling competence and patterning windows

We also uncovered that cell density regulates signaling competence, which thereby influences the timing of the WNT response window. Specifically, the ability of cells to respond to WNT and undergo mesoderm differentiation depended on seeding density. Even in unpatterned cells at too low or too high a density an otherwise optimal WNT stimulus failed to rescue mesoderm differentiation. Thus, we concluded that cell density sets the timing of the WNT competence window. These findings are consistent with previous discoveries that the BMP-driven mesoderm induction only occurs above a critical cell density, failing entirely at sparse cell seeding^5,32^. Additionally, we and others have noted how cell density influences the spatial pattern of germ layer differentiation, with more crowded colonies tending to form a narrower mesoderm band^13^.

These observations indicate that cell density, either through mechanical forces, the number of neighbors, or perhaps even the aspect ratio of polarized cells fundamentally tunes how cells decode signaling dynamics. This interplay between density and timing means that what might appear as an autonomous “developmental clock” is actually tunable based on cell-cell proximity and communication. Thus, any rules for temporal morphogens must likely account for the local cell density. Intriguingly, *in toto* imaging of mouse gastrulation has revealed that the variance in cell density is locally minimized at the site of PS formation^50^. We surmise that the interaction between cell density and dynamic morphogen decoding could serve as a mechanism to ensure tissues form only after they have achieved the correct size or number of cells.

### Optogenetic activation can enhance pattern fidelity beyond natural mechanisms

A technical innovation was the use of optogenetic activation of WNT signaling, which allowed us to control this morphogen with unprecedented precision in space and time. This approach contrasts with traditional ligand-based stimulation and proved to have distinct advantages in both dissecting pattern formation and engineering fate boundaries. First, optogenetic manipulation of the temporal features of WNT signaling allowed us to avoid complications with washing Wnt ligand (which is not practical due to its palmitoylated residues^51,52^) and, by leveraging the LITOS 96-well light delivery plate, allowed us to practically screen through a much larger space of signaling dynamics than have ever been previously explored. Second, using DMDs to project a spatial pattern of WNT we were able to create a sharply defined boundary between mesoderm and adjacent fates. This precision was striking as the mesoderm domain did not fade gradually fade into neighboring fates as occurs when patterning solely relies on diffusion-based waves of endogenous morphogens. Thus, optogenetic control produced boundaries that were sharper than those generated by exogenous ligand, highlighting that bypassing endogenous transport and feedback can uncouple pattern fidelity from the constraints of natural signaling architectures. Alternatively, this boundary region of mixed fates could generate important intermediate cell types. Indeed, notochord progenitors and neuromesodermal progenitors (NMPs) for example coexpress SOX2 and BRA^53–56^. Future experiments utilizing spatial transcriptomics in the 2D gastruloid model could help clarify the developmental significance and potential functional roles of these intermediate boundary regions.

### Implications for developmental biology and regenerative medicine

These findings illustrate the principles of the role of timing and dynamics in morphogen pathways encoding developmental information. They reinforce the emerging view that cells of the embryo are not simply reading static morphogen concentrations, but are in fact decoding the history of signal exposure. This parallels other observations in the 2D gastruloid such as the finding that the time-integral of BMP signaling (rather than the instantaneous level) determines the fate of the amnion^18^ and that cells respond to the time derivative of NODAL signaling^57^. Furthermore, we show that by altering the timing of WNT signaling alone, total inversion of the germ layers occurs, with mesoderm peaking at the center of the colony as opposed to the outside edge. The observations of the importance of sequential and precisely timed delivery of morphogens underscore that developmental timing is an essential fourth dimension of pattern formation.

Our findings also have practical implications for organoid models and stem cell-based regenerative medicine therapies. Creating stable and sharply bounded tissue domains remains an ongoing challenge in the field of regenerative medicine^58–61^. Engineering organoids that achieve correct spatial organization and cell type proportions has remained a challenge. Our insights into the importance of timing suggest that delivering signals in controlled pulses or sequences (rather than continuous exposure) may better mimic the embryos natural patterning cues and thus coax cells into a more accurate organization. It also suggests that subtle refinements in the relative timing of signals could be a controlled method to tune the ratios of cell types. Current mesoderm subtype differentiation protocols don’t take the precise timing of signals into consideration^47,62,63^. Ultimately, defining the “rules” of tissue patterning, such as the WNT/BMP timing rules defined here, will enhance our ability to build organized tissues and perhaps even entire embryonic structures in a dish, opening new avenues in developmental biology, disease modeling, and regenerative medicine.

### Limitations and Future Directions

While our study defines spatiotemporal rules for germ layer patterning in the 2D gastruloid, it is important to recognize the system’s limitations. Our 2D monolayer system likely lacks the mechanical, geometric, and extraembryonic interactions present in the developing embryo, limiting the recapitulation of full morphogenetic behaviors such as primitive streak elongation and cellular ingression. Future experiments using more complex 3D gastruloids or embryo-like models in combination with 2-photon excitation of optogenetic tools will be essential to determine whether the temporal rules we identified persist in 3D where tissue geometry can feedback into signaling and cell fate choices. Furthermore, our focus was primarily on the WNT and BMP pathways, without explicit control over other crucial signaling networks involved in gastrulation, such as FGF and retinoic acid. Similar studies leveraging ERK KTRs (Kinase Translocation Reporters) and opto-FGF or opto-SOS tools could similarly dissect the decoding function of those pathways into cell fates. In fact, multiplexed use of red, green, and blue-light responsive optogenetic tools targeting multiple pathways promises to more fully unravel the signaling logic underlying gastrulation. Finally, our scRNA-seq analysis suggested distinct mesoderm subpopulations emerge based on WNT timing; future targeted experiments examining lineage-specific markers will be crucial for understanding the molecular drivers behind these distinct fates. In summary, these findings offer fundamental insights into the importance of signaling dynamics in developmental systems and lay groundwork for future efforts in developmental biology, gastruloid modeling, and regenerative medicine, where precise temporal control of signaling emerges as a critical factor for engineering complex tissues and organs.

## Resource Availability

### Lead Contact

Further information and requests for resources and reagents should be directed to and will be fulfilled by the lead contact, Maxwell Wilson.

### Materials Availability

Plasmids generated in this study are available upon request to Maxwell Wilson without restrictions.

### Data and Code Availability

All code generated during this study has been deposited at the Wilson Github repository. Any additional information required to reanalyze the data reported in this paper is available from the lead contact upon request.

## Acknowledgements

This work and M.Z.W. was supported by R01 HD108803-01. N.B. was supported by NIH F31HD108952-01, J.R. was supported by NIH F32HD114539-01, and R.P. was supported by the CIRM Research Training Grant EDUC4-12821. S.S.D was supported by NIH R01HD099517, NIH R01HG011013 and NSF 2339849 grants. Some schematics in this manuscript were created using BioRender.com. We would also like to thank Eric Siggia for his constructive feedback in drafting the manuscript, and Dennis Clegg for being a co-sponsor of N.B.

## Author contributions

Conceptualization, N.B. and M.Z.W.; methodology, N.B., R.P., and M.Z.W.; software and data analysis, N.B., J.R., and M.Z.W.; investigation, N.B., and M.Z.W.; writing – original draft, N.B. and M.Z.W.; editing –S.S.D. and J.R.; final draft N.B. and M.Z.W.; funding acquisition N.B. and M.Z.W.

## Declaration of interests

M.Z.W. is an employee, shareholder, and board member of Integrated Biosciences Inc..

## Supplemental Figures

**Figure S1.**
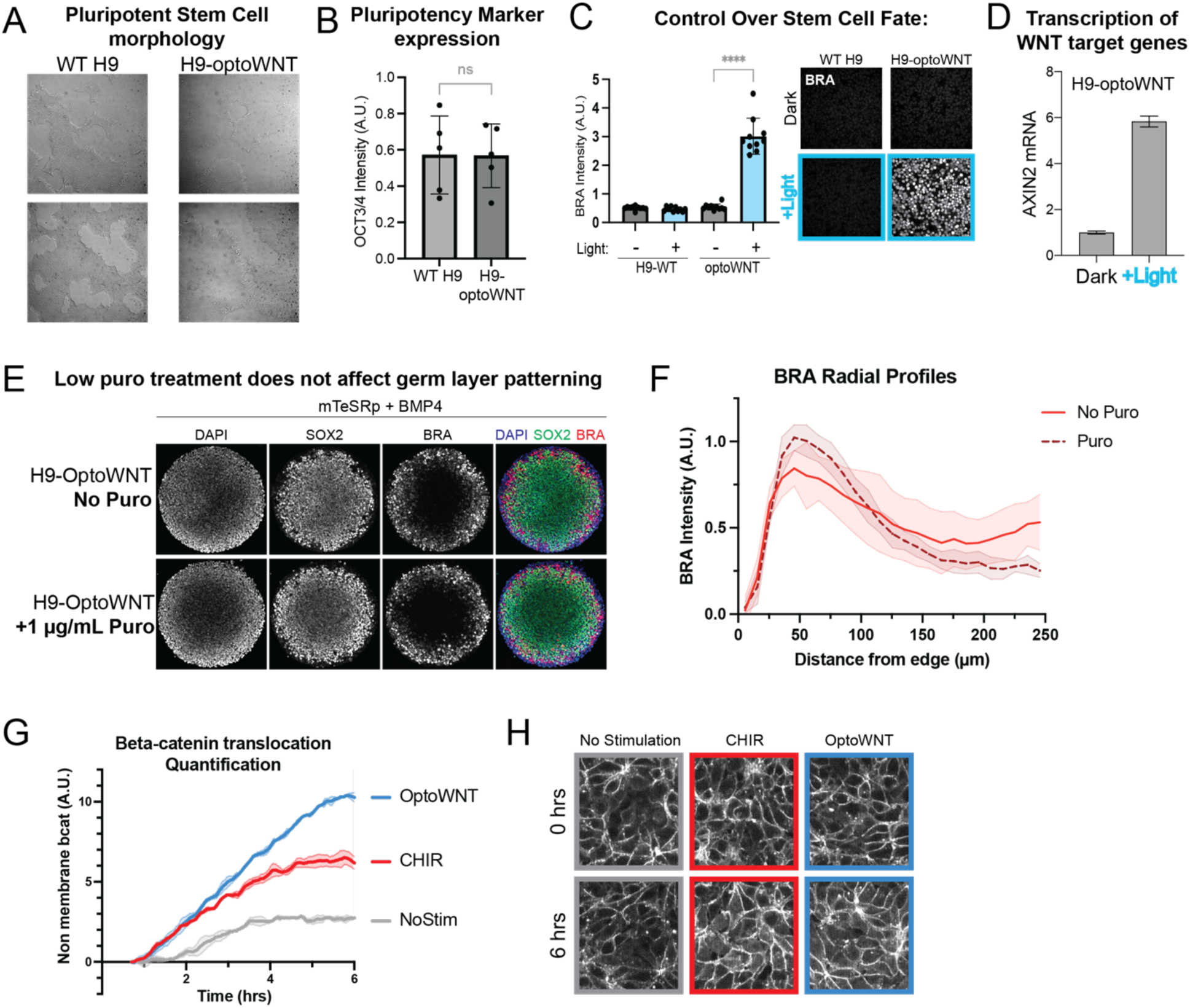
H9-OptoWNT cell line retains pluripotency compared to WT-H9 hESCs and light activation mimics canonical WNT signaling. (Related to figure 1) **(A)** Brightfield images of WT H9 cells compared to H9-optoWNT cells. **(B)** OCT3/4 expression detected and quantified via immunofluorescence analysis in WT H9 cells compared to H9-optoWNT cells. **(C)** BRA differentiation in response to blue light stimulation detected via immunofluorescence analysis. **(D)** RT-qPCR of Axin2 mRNA levels in H9-optoWNT cells in response to light stimulation for 24 hours. **(E)** Representative IF stained colonies for H9-optoWNT cells treated with and without puromycin during germ layer formation. **(F)** BRA radial profiles for 5 representative colonies with and without puro treatment, quantified via DAPI nuclear segmentation. (G-H: beta catenin translocation for OptoWNT stimulation compared to CHIR 99021). **(G)** Mean and standard deviation for the centered rolling average of cells treated with 6 hrs of CHIR (red) and OptoWNT (blue) compared to no stimulation (grey). **(H)** Representative fields of view of beta catenin-tdmruby before (top) and after (bottom) 6 hrs of stimulation, no stimulation (*left*), stimulation with CHIR (*middle*), and OptoWNT (*right*).

**Figure S2.**
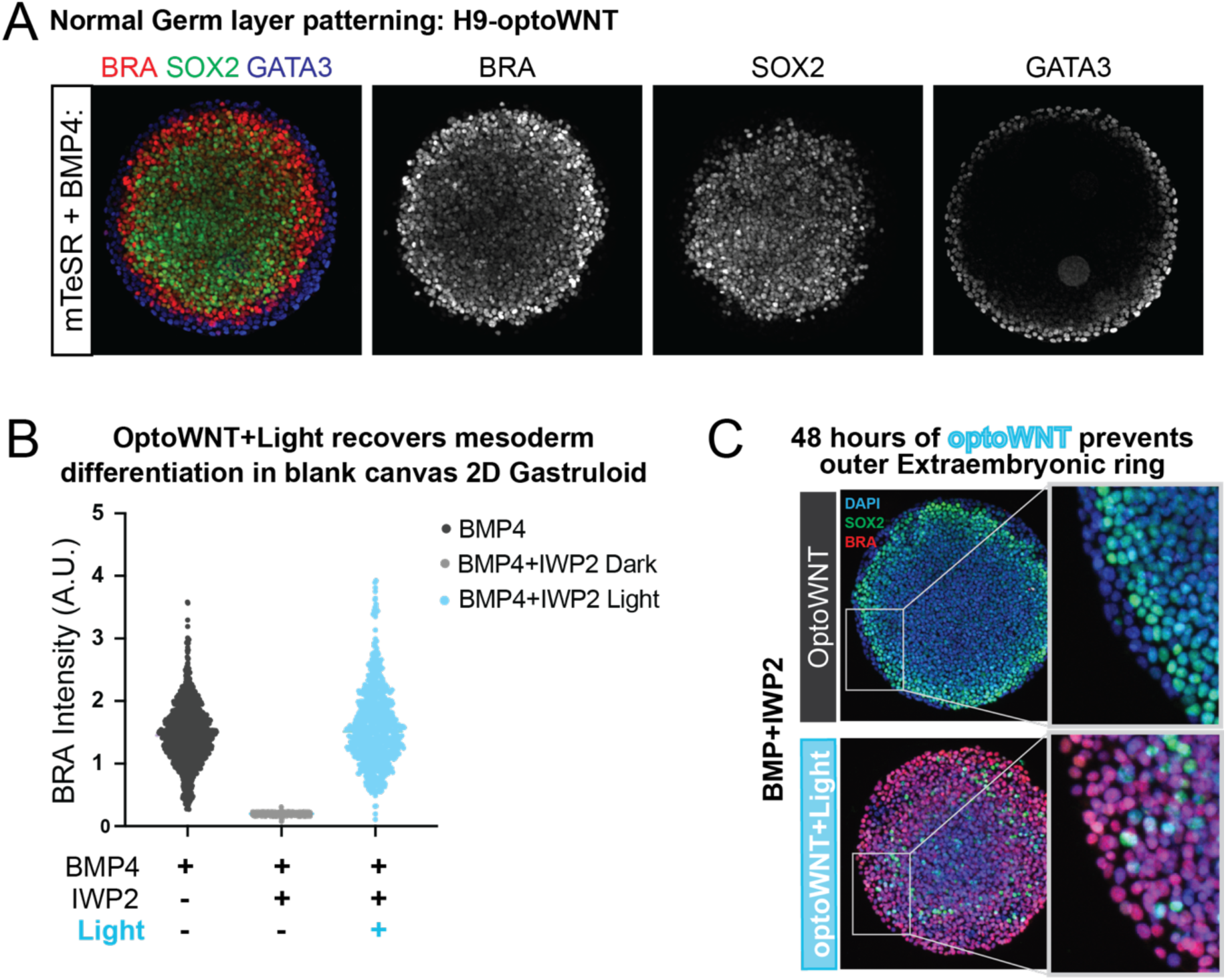
Optogenetic WNT activation recovers mesoderm differentiation but prevents in “blank canvas” 2D gastruloid (Related to figure 1) **(A)** Individual channels for micropatterned H9-optoWNT treated with BMP4 for 48 hours and stained for the germ layers (colony shown in fig 1C) **(B)** Quantification of Brachyury in blank canvas experiment compared to BMP4 treated control colonies for 3 replicate colonies. BRA intensity quantified via DAPI nuclear segmentation. **(C)** H9-optoWNT colonies treated with BMP and IWP, IF stained for DAPI and indicated fate markers. Boxes highlighting that OptoWNT stimulation for 48 hours prevents differentiation of the outer extraembryonic ring as indicated here by DAPI, SOX2-. (colonies shown in Fig 1I)

**Figure S3.**
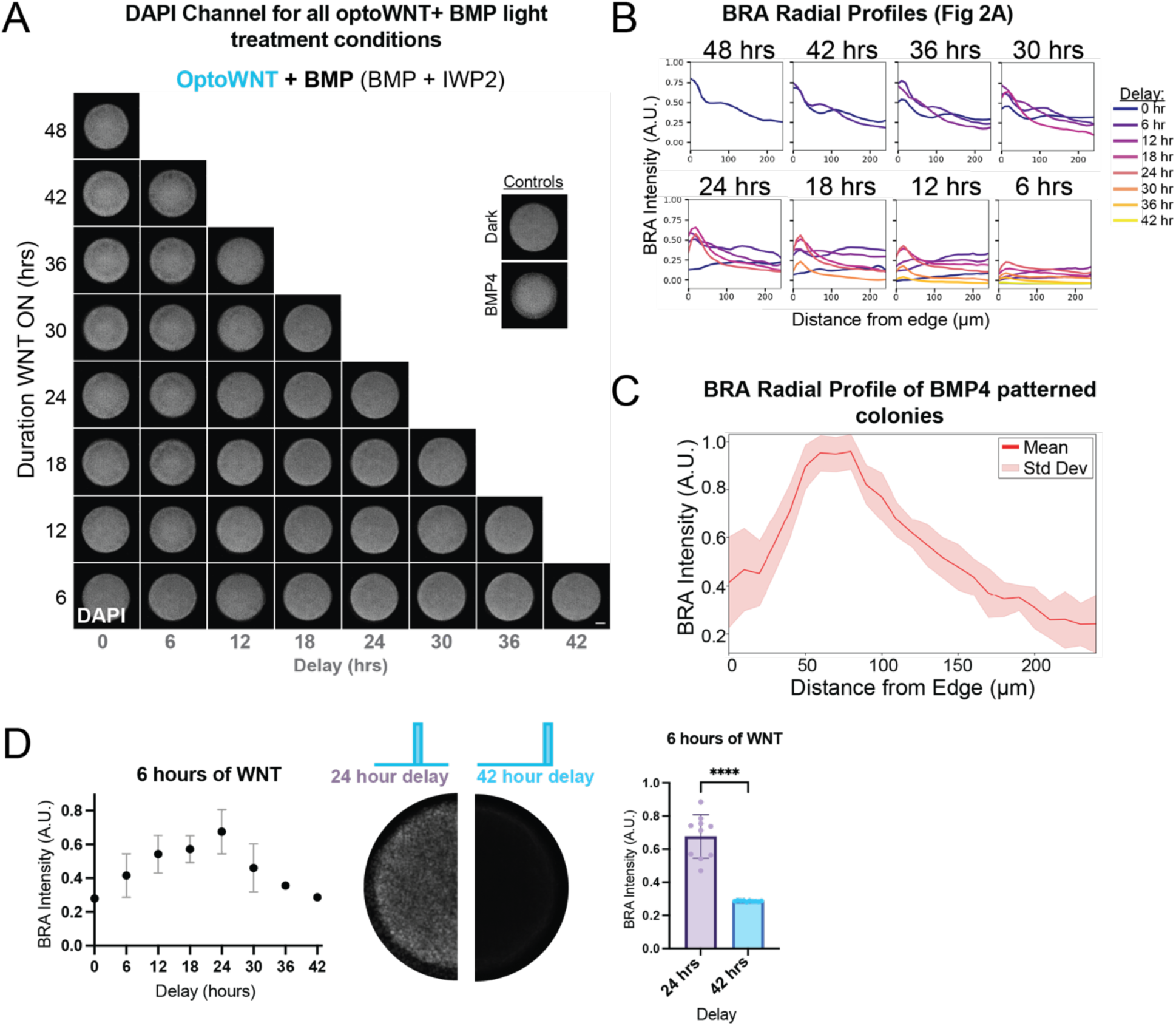
The timing of OptoWNT modulates mesoderm differentiation in Gastruloids treated with BMP+IWP2. (Related to Figure 2) **(A)** Colony averages of DAPI channel for all colonies shown in Fig 2A. 10 replicate colonies per treatment. **(B)** BRA radial profiles for each light treatment in Fig 2A. **(C)** BRA radial profile of BMP4 patterned colonies (10 replicate colonies). **(D)** Representative light treatments demonstrating that the timing of the WNT signal affects the amount of mesoderm differentiation for 6 hours of OptoWNT. *Left*: BRA differentiation for 6 hours of WNT at each delay indicated. *Middle*: BRA colony averages with 24- and 42-hour delays. *Right*: differences in BRA differentiation at 24 and 42 hour delays is statistically significant ****P<0.0001

**Figure S4.**
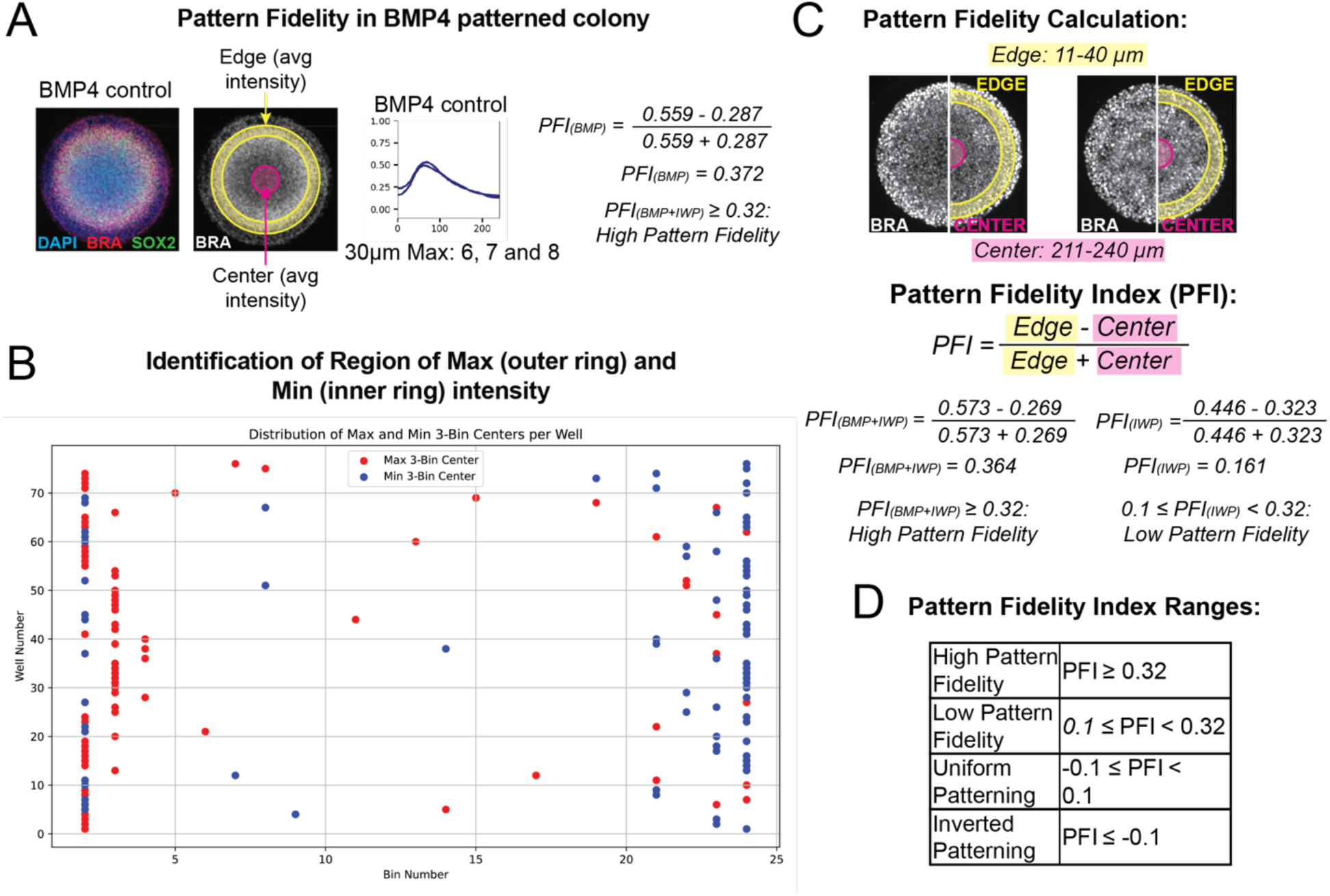
The Pattern Fidelity index is a metric that reflects the expected difference in mesoderm differentiation at the edge compared to the center of the colony. **(A)** In a BMP4 patterned 2D gastruloid, Brachyury patterns in a ring that is a few cell widths in from the edge of the colony. The region of max BRA intensity is in this ring (yellow), and the BRA signal in the center of the colony should be much lower (magenta). Calculation (*left*): PFI for BMP4 patterned colonies results in high pattern fidelity. **(B)** Visualization of the 3-bin max (red dots) and 3-bin min (blue dots) for each light treatment to determine the “edge” and “center” regions for optogenetically patterned colonies. **(C)** Pattern fidelity calculations for the optoWNT patterned colonies shown in Fig 3C. These gastruloids treated with OptoWNT+BMP (*left*) result in “high” pattern fidelity, while those treated with OptoWNT alone (*right*) result in low pattern fidelity.

**Figure S5.**
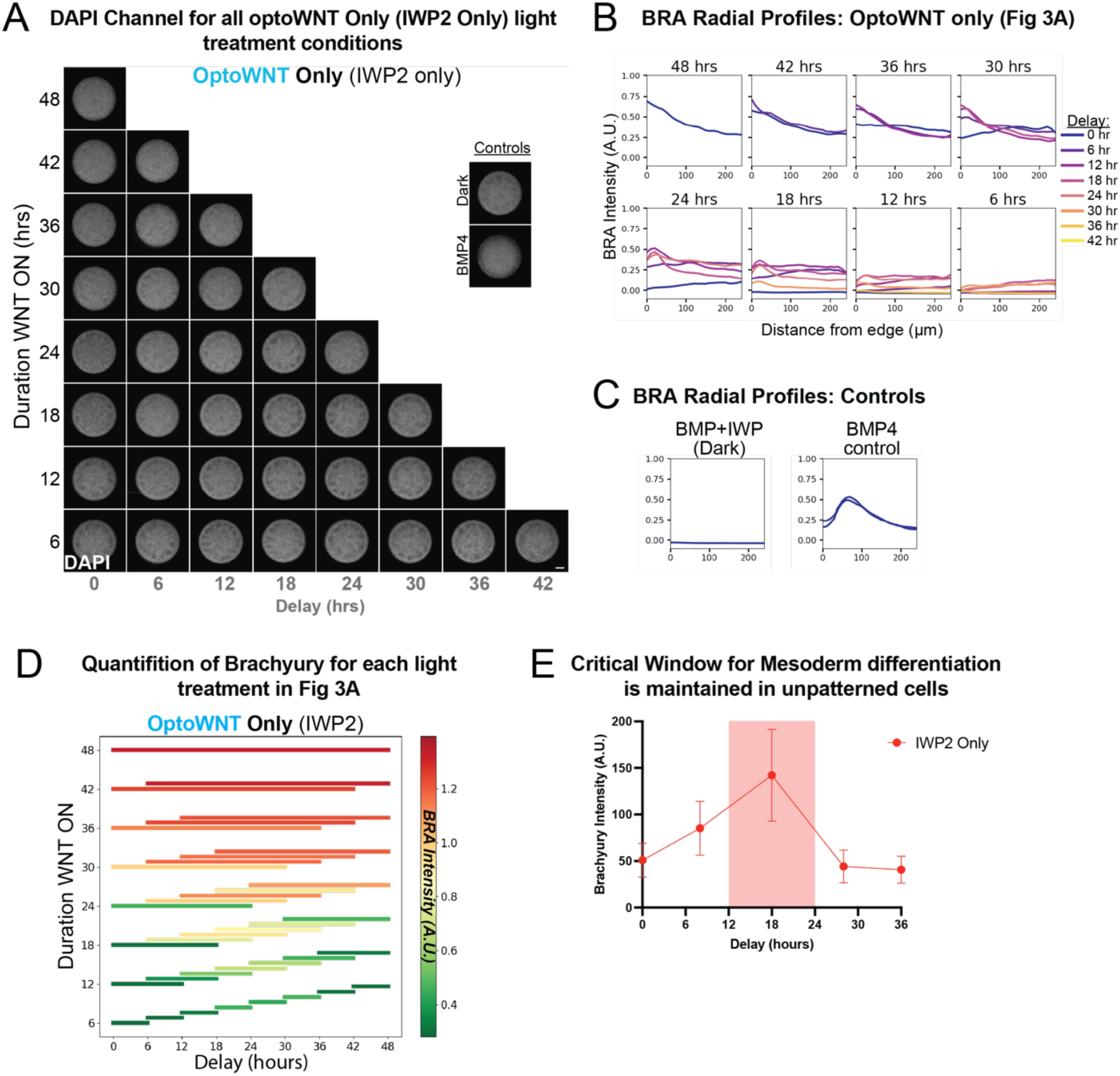
The critical window for mesoderm differentiation exists without BMP4 and in monolayer culture. (Related to Figure 3) **(A)** Colony averages of DAPI channel for all colonies shown in Fig 3A. 10 replicate colonies per treatment. **(B)** BRA radial profiles for each light treatment in Fig 3A. **(C)** BRA radial profile of control patterned colonies (10 replicate colonies per line). **(D)** Heat map bar plot representing the average mesoderm differentiation for each light treatment over 10 replicate colonies shown in 3A. The color of each bar represents the intensity of Brachyury (red is the highest intensity and green is the lowest). The length and location of each bar represents the duration and timing of the WNT signal respectively. The mean level of mesoderm differentiation per light treatment was quantified based on DAPI nuclear segmentation. **(E)** Quantification of Brachyury for H9-optoWNT cells plated in monolayer (not micropatterned) culture, and treated with 12 hrs of OptoWNT at the indicated delays. Detected via IF and DAPI nuclear segmentation.

**Figure S6.**
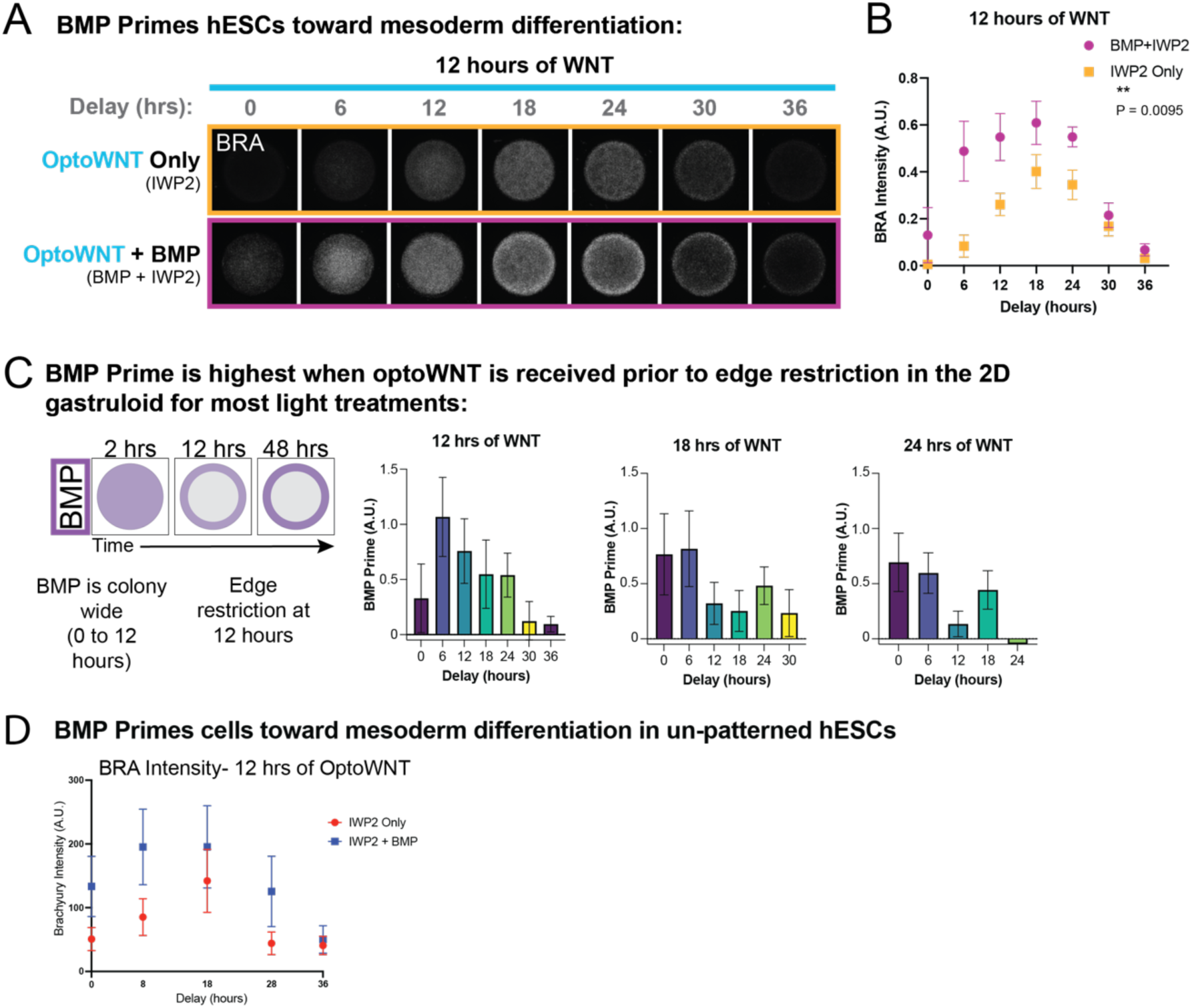
BMP primes cells toward mesoderm differentiation through some other mechanism besides WNT ligand secretion. (Related to Figure 3) **(A)** Comparison of gastruloids treated with 12 hours of blank canvas OptoWNT at different delays, with and without BMP, demonstrating the increase in mesoderm differentiation when BMP4 is present despite receiving the same duration of WNT and WNT ligand secretion inhibited. Colony average of 10 replicate colonies. **(B)** Quantification of Brachyury for the colonies shown in (A). (C) *Left*: schematic demonstrating the timing of BMP signaling edge restriction that occurs at around 12 hours. The BMP prime is generally more pronounce when the WNT signal is received prior to edge restriction. **(D)** BMP primes cells toward WNT driven mesoderm differentiation in unpatterned cells. H9-optoWNT cells plated in monolayer (not micropatterned) culture, and stimulated with 12 hrs of OptoWNT at the indicated delays, treated with optoWNT Only (IWP2 only, red) and OptoWNT + BMP (IWP2+BMP, blue).

**Figure S7.**
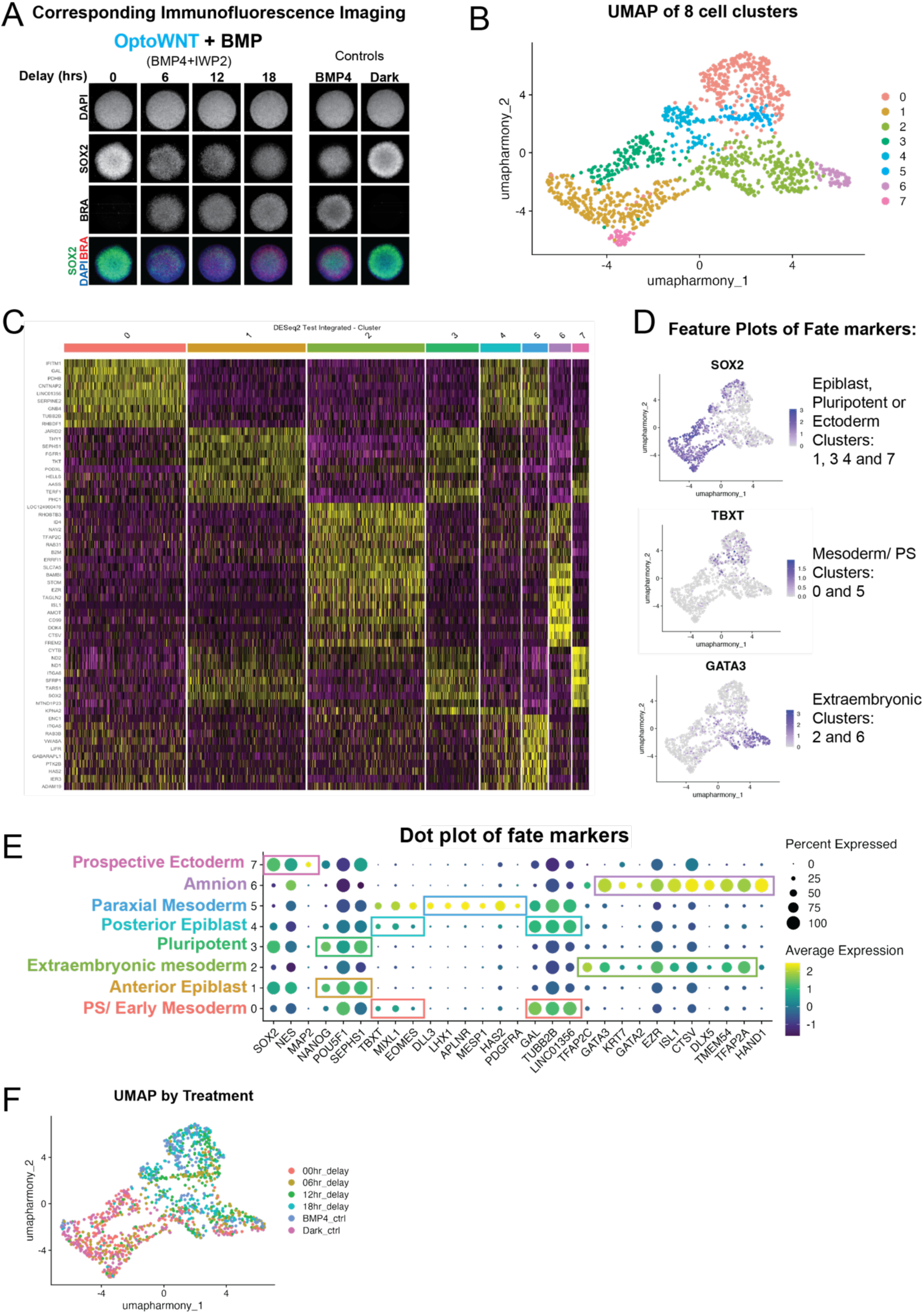
Corresponding Immunofluorescence analysis and scRNAseq cluster cell type annotation. (Related to figure 4) **(A)** Corresponding immunofluorescence analysis for each treatment included in the scRNAseq experiment. Colony averages of 10 replicate colonies. The same LUTS were used for all conditions. **(B)** The 8 cell clusters identified via unsupervised clustering and corresponding cluster numbers. **(C)** Differential gene expression across the 8 cell clusters. **(D)** Feature plots of fate markers identified clusters as extraembryonic (clusters 2 and 6), mesoderm or primitive streak lineage (clusters 0 and 5), or epiblast/ pluripotent/ ectoderm (clusters 1, 3, 4, and 7). **(E)** Dot plot of key marker genes to further elucidate the identity of each cluster. **(F)** UMAP colored by treatment.

**Figure S8.**
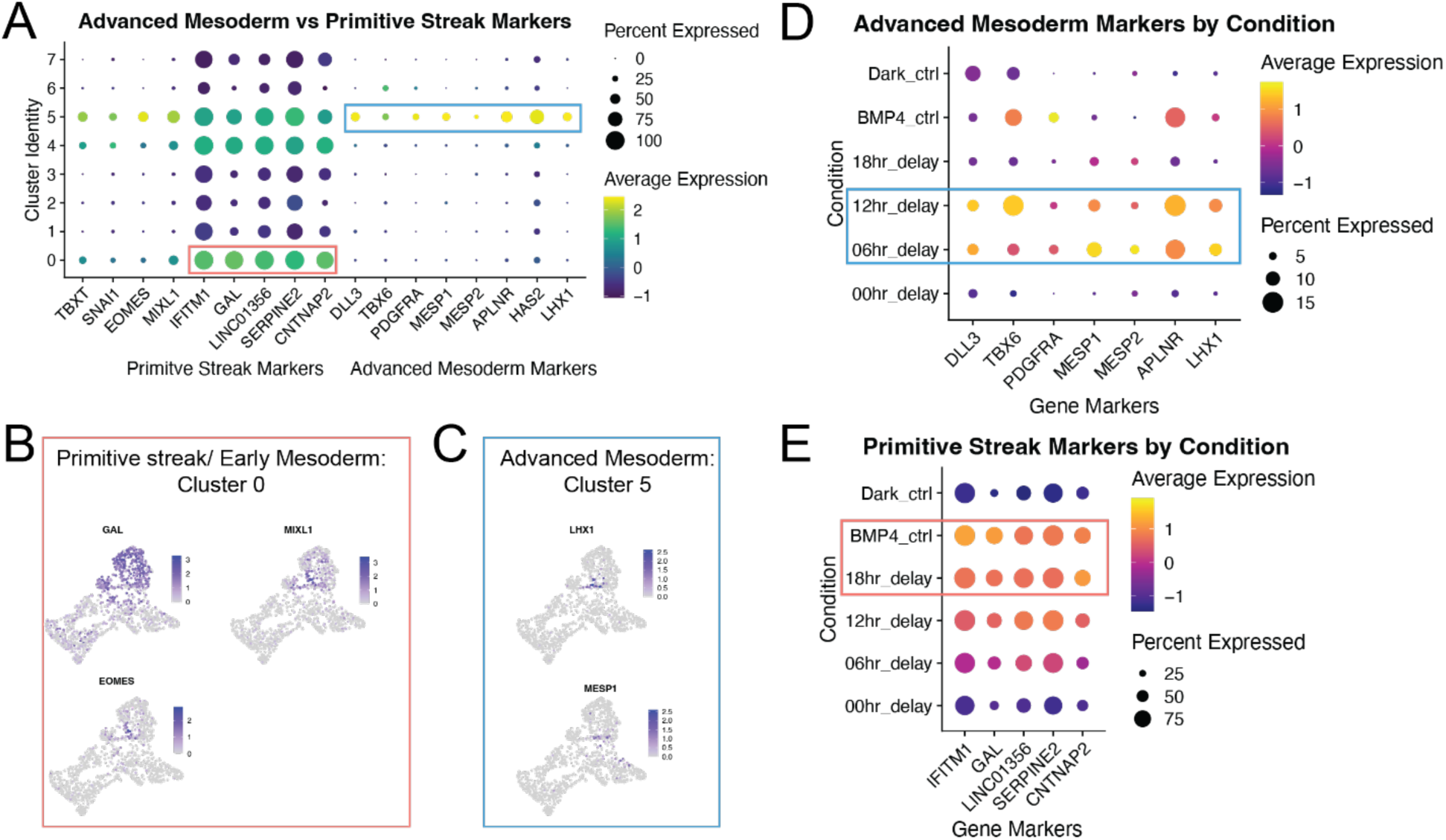
OptoWNT patterned 2D gastruloids with 6- and 12- hour delays differentiate cells into advanced mesoderm subtypes. (Related to figure 4) **(A)** Dot plot of advanced mesoderm and primitive streak markers distinguishing cluster 5 as a more advanced paraxial mesoderm population and cluster 0 as an early mesoderm/ primitive streak population. **(B)** Feature plots of primitive streak and early mesoderm marker genes. **(C)** Feature plots of more advanced paraxial and intermediate mesoderm markers. **(D)** Dot plot of advanced mesoderm markers by condition. The 12- and 6-hour delays (aberrant patterning phenotypes) result in high expression of advanced (paraxial, lateral plate and intermediate) mesoderm subtypes (percent is of all cells from that treatment, not cluster). **(E)** Dot plot of primitive streak/ early mesoderm markers by condition. BMP4 (control) and partial rescue (12 hrs optoWNT with 18-hour delay) patterned colonies express higher levels of primitive streak marker genes. These genes are expressed at much lower levels in the aberrant patterning phenotypes (0-, 6-, and 12-hour delays).

**Figure S9.**
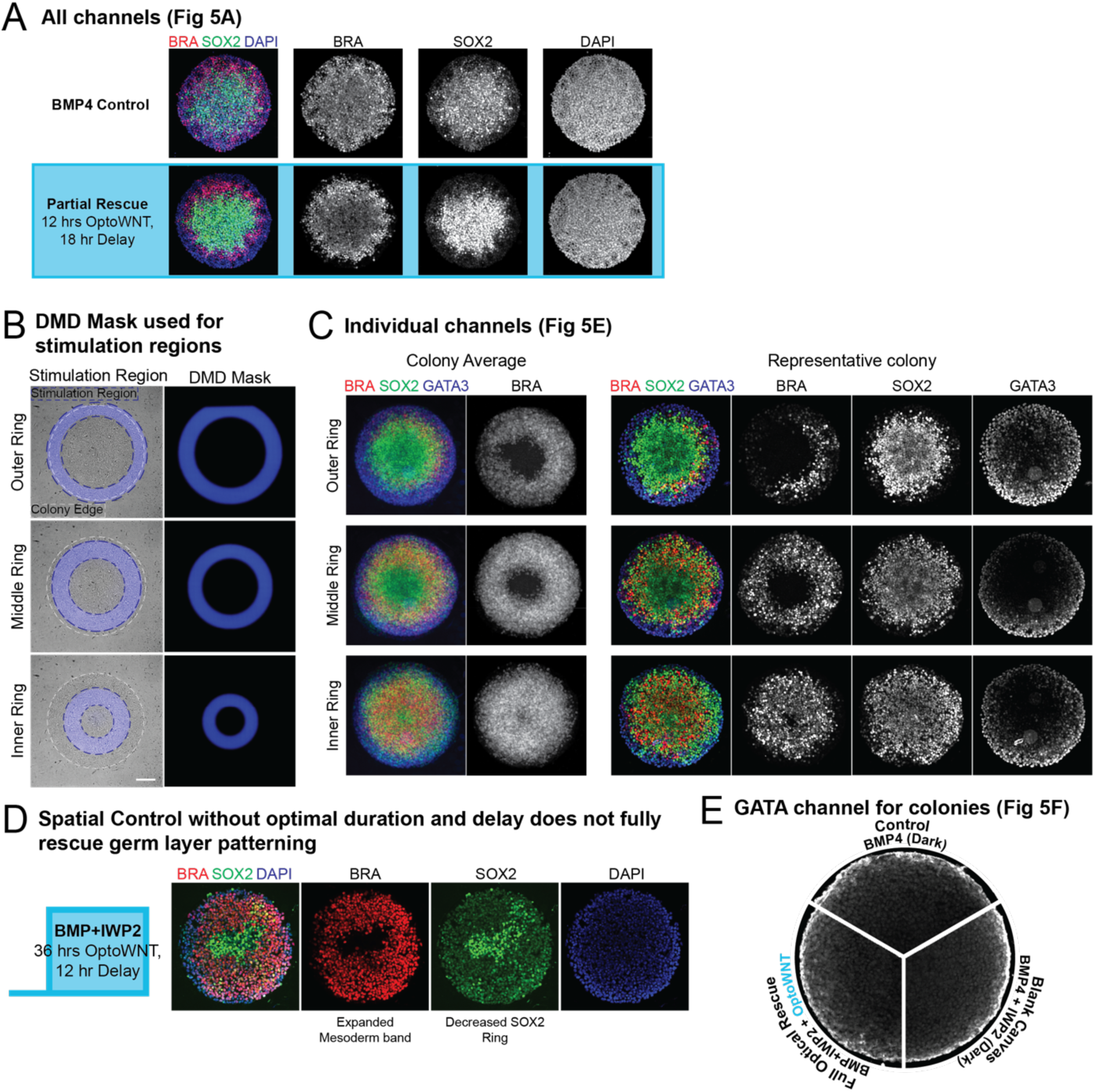
Spatiotemporal control of OptoWNT rescues BRA and GATA3 patterning. (Related to figure 5) **(A)** All channels shown in antibody-stained colony in figure 5A highlighting the differences in 2D gastruloid boundary formation in a normally patterned BMP4 colony compared to the partial optical rescue. **(B)** The DMD masks used for each stimulation region. **(C)** Additional channels for the colony average and representative colonies shown in 5E. **(D)** Gastruloids patterned with spatial control (middle ring stimulation region) but stimulated with a non-optimal duration and delay results a mesoderm band that is too wide and a decreased SOX2+ inner ring. **(E)** GATA3 channel for the colonies shown in figure 5F.

## Supplementary Notes

### Supplementary Note 1: Annotation of cell types in scRNAseq clusters

Unannotated clusters (also shown in **fig S7B**):

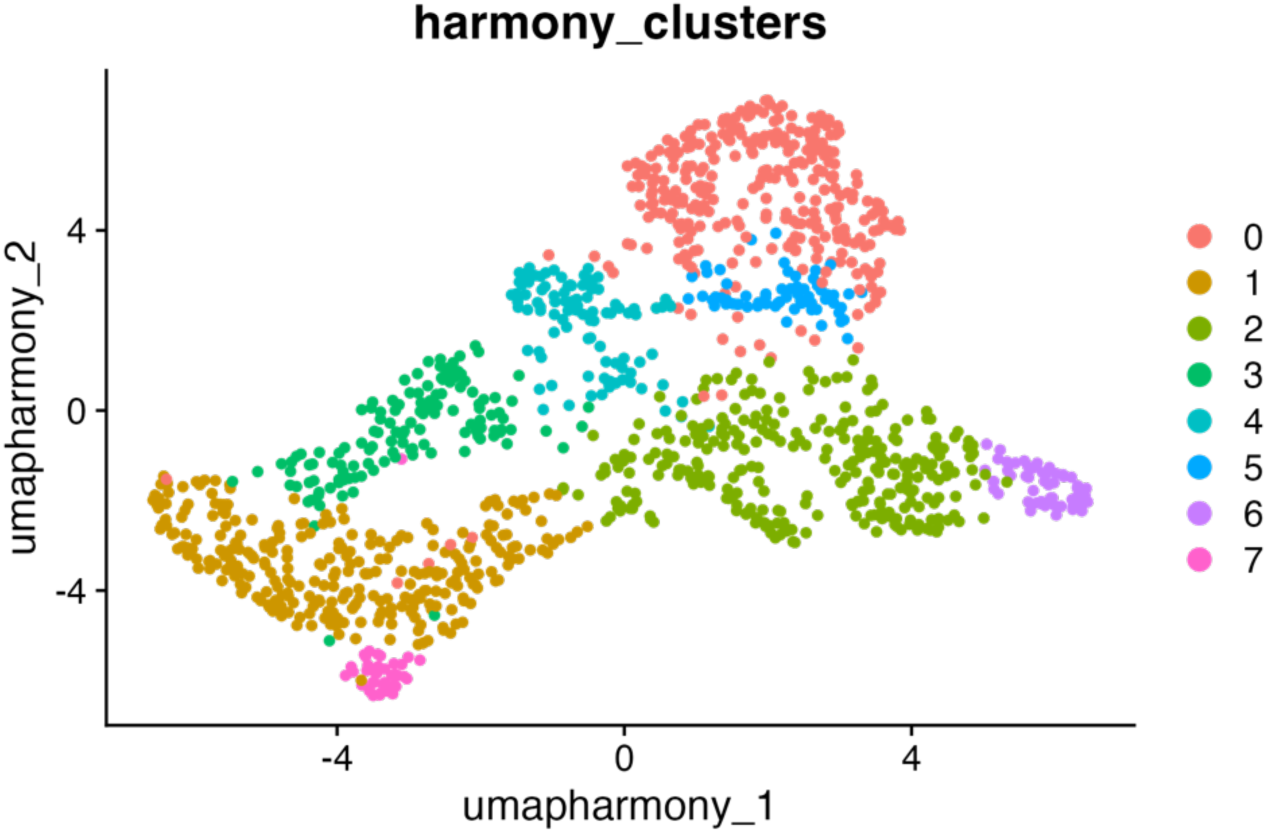

There were 2 clusters, cluster 0 and cluster 5 (**fig S7B**), that clearly corresponded to mesodermal lineages as indicated by expression of TBXT (**fig S7E**). The cluster that we identified as primitive streak/ early mesoderm (cluster 0) is enriched for canonical primitive streak markers (TBXT, MIXL1, EOMES) and genes that are highly expressed in primitive streak and midline mesoderm in a CS7 embryo (GAL, SERPINE2, CNTNAP2)^38^ (**fig S7E**). There is some expression of SOX2 in this cluster while the other mesodermal population (cluster 5) has negligible SOX2 expression (**fig S7D**) supporting a primitive streak identity rather than a committed advanced mesoderm population. Therefore, we identify this population as primitive streak and possibly primed for midline mesoderm fates such as the notochord and axial mesoderm. Co-expression of both primitive streak and mesoderm markers led to the identification of PS/early mesoderm representing the forming mesoderm.

The cluster identified as Paraxial and lateral plate mesoderm (cluster 5) is strongly enriched for a mesodermal population that has already traversed the primitive streak (LIFR, ENC1, APLNR, LHX1, MESP1)^12^ and is highly migratory (RAB3B, PTK2B, ITGA5) (**Supp Table 3**). DLL3, TBX6, MESP1 and MESP2 are paraxial mesoderm markers^47^ that are all expressed in this cluster. There is also some expression of lateral plate markers including APLNR^64^, suggesting this cluster could contain subclusters of both paraxial and lateral plate mesoderm. However, the paraxial signature is stronger (fig S8D). This cluster also has much lower expression of pluripotency markers (SOX2, POU5F1 and SEPHS1) (**fig S7E**) and highly expresses genes involved in ECM remodeling and integrin signaling which are hallmarks of EMT^65^ further supporting the paraxial/ lateral plate mesoderm identity.

The genes expressed in two of the clusters were consistent with extraembryonic identities (GATA3, ISL1, and EZR). In one of the clusters, HAND1 is specifically and significantly upregulated supporting an amnion identity^66^. Compared to the amnion cluster, expression of extraembryonic specific markers is lower in the other cluster, and the top differentially expressed genes suggest a high BMP signature (BMP4, BAMB, ID4), and reveal some ECM and motility genes characteristic of extraembryonic mesoderm (HPGD, SEM6D)^67,68^. Extraembryonic mesoderm is in contact with amnion in a gastrulating human embryo, and was recently identified in 2D gastruloids^39,40,69^, leading to the identification of the other extraembryonic lineage as extraembryonic mesoderm.

We found three clusters that highly express SOX2 (Clusters 7, 3 and 1: **fig S7E**). The gene expression patterns of these three clusters suggest they all retain pluripotency rather than representing specified ectoderm. Cluster 7 highly expresses MAP2^70^, SFRP1^71^ and NES^72^ and has lower pluripotency marker expression (POU5F1, NANOG, SEPHS1) than clusters 3 and 1, suggesting neuro-ectoderm. However, the pluripotent and metabolically active signature of the top differentially expressed genes (**Supp table 3**) suggests this population is not yet committed to ectodermal fates leading to the *prospective* neuro-ectoderm identification.

Cluster 3 highly expresses SOX2 and other pluripotency markers but does not significantly upregulate any posterior or anterior epiblast identity genes and exhibits lower expression of neuroectoderm marker NES than cluster 1. The top differentially expressed genes in this cluster (**Supp table 3**) contain pluripotent and proliferative genes suggesting a pluripotent epiblast population not primed for either anterior or posterior fates. Additionally, cells in this cluster are only present in the dark (no WNT) and 0 hr delay (no mesoderm) treatment conditions (**Fig 4E**), and are not present in BMP4 patterned colonies, further supporting the pluripotent identity of this cluster.

Cluster 1 highly expresses pluripotency and epiblast markers (SOX2, NANOG, POU5F1, and SEPHS1), in addition cell cycle genes (**Supp table 3**). It also expresses neuroectoderm markers such as NES suggesting this population has not been exposed to posteriorizing cues, and leading to the identification of epiblast primed for anterior fates identity. SOX2 expression is also upregulated in Cluster 4 (**fig S7E**), and this cluster expresses primitive streak markers TBXT, MIXL1, EOMES, GAL, and TUBB2B at lower levels than the mesodermal populations. The top differentially expressed genes (**Supp table 3**) are suggestive of a highly proliferative and more pluripotent population than clusters 0 and 5, leading to the identification of epiblast primed to ingress through the primitive streak, or “posterior primed epiblast”.

## METHOD DETAILS

### Cell lines

All experiments were conducted using H9 human embryonic stem cell lines (WA09) (provided by the UCSB Stem Cell Facility). A clonal β-catenin reporter line was generated through CRISPR/Cas9-mediated homology directed repair. The piggyBAC transposase-based insertion system was used to stably integrate the optogenetic tool for controlling WNT into the H9 hESC tdm cell line (H9-OptoWNT). Pluripotency was confirmed by colony morphology, staining of pluripotency markers OCT3/4 and SOX2, and confirming unaltered differentiation into the three germ layers as compared to unedited H9 hESCs. All cell lines routinely tested negative for mycoplasma contamination.

### Cell Culture

H9 hESCs were grown and maintained in mTeSR Plus feeder-free maintenance medium (StemCell Technologies, Cat No. 100-0276) and cultured on Matrigel hESC-Qualified Matrix, LDEV-free (Corning, Cat No. 354277) coated dishes for routine maintenance. Matrigel dishes were coated overnight at 4°C and incubated at room temperature for at least one hour prior to seeding. Cells were passaged every 4-6 days as cells reached 80% confluency using ReLeSR (StemCell Technologies, Catalog # 100-0483). All experiments were completed within 3 to 10 passages after thaw of the working bank.

### Generation of the H9-hESC OptoWNT CRISPR tagged b-catenin tdmruby cell line

Single cell H9 hESCs were seeded onto Matrigel coated plates and transfected with Lipofectamine STEM transfection reagent (Invitrogen, Cat No. STEM00015) according to manufacturer’s instructions. Transposase and transposon (pPig-oLRP6 plasmid) were mixed into Lipofectamine STEM transfection complexes in a ratio of 1:3 (w/w). Transfected cells were expanded then selected with 2 µg/mL puromycin, and FACS sorted on a SH800 Sony Cell Sorter to isolate clonal populations. Clonal populations were screened for pluripotency their ability to recapitulate canonical WNT signaling in response to light. To ensure a high optoWNT transgene expressing population in culture, OptoWNT H9s were treated with 1 µg/mL puromycin for 24 hours the day after each routine passage. This is a frequently utilized approach to maintain stable transgene expression in hESCs and had no effect on PSC morphology and marker expression over many passages. To maintain stable expression of the transgene during 48-hour differentiation experiments, we tested a range of concentrations of puro (0.5 µg/mL, 1µg/mL, 2 µg/mL, and 4 µg/mL) during differentiation into the three germ layers. These concentrations had no effect on normal germ layer patterning (**fig S1E,F**), so we chose the lowest concentration the efficiently maintained expression of the transgene (1 µg/mL).

### Micropatterning and differentiation experiments

Micropatterning experiments were completed using 96 well glass bottom plates with 500 µm diameter micropatterned discs purchased from CYTOO. Coating was done per manufacturer’s instructions. Briefly, wells were incubated with 10 µg/mL CellAdhere™ Laminin-521 Matrix (StemCell Technologies, Catalog #77003) diluted in PBS for 2-3 hours at room temperature. Wells were rinsed by manually adding and removing PBS using a micropipette to ensure 50 µL of liquid was left in the wells at all times. The washing step was repeated three times prior to plating cells. Cells were dissociated into single cells with Accutase™ (StemCell Technologies, Catalog #07920) and seeded at 50,000 cells per well (1,500 cells/mm^2^) unless specified otherwise. H9-OptoWNT cells were seeded in 1 µg/mL puro to ensure stable transgene expression for optogenetic experiments. Differentiation experiments were completed after overnight incubation. BMP4 patterned colonies were treated with 50 ng/mL Human Recombinant BMP4 (StemCell Technologies, Catalog #78211) diluted in mTeSR plus. Colonies patterned with OptoWNT were treated with 50 ng/mL Human Recombinant BMP4 and 2 µM IWP-2 (Selleckchem) to create the blank canvas of WNT signaling.

### Optogenetic Stimulation

Temporal light patterns were controlled using a LITOS (LED Illumination Tool for Optogenetic Stimulation) device. Cells seeded in 96 well plates were placed onto a LITOS device kept in a standard tissue culture incubator at 37°C and 5% CO2 for the duration of the experiment. User defined illumination patterns were uploaded to the LITOS device to achieve independent control of illumination for each well. Optogenetic stimulation was achieved with blue LED light (465–470 nm) at max power density (135 ± 0.11 µW/cm^2^) and activated for 1 second every 20 seconds.

Spatial patterning of light was achieved using digital micromirror devices (DMDs) integrated into our microscope’s optical path triggered via Nikon NIS Elements software. NIS Elements software was used to define the ROI for stimulation and a Jobs definition was used to define stimulation parameters and timing. The stimulation parameters were 1s pulses every 60s at 25% LED power (λ = 455nm). All optogenetic experiments were optimized to maintain continuous (maximal) activation of CRY-2 during activation windows without phototoxicity so the method of light delivery would not alter the degree of optogenetic activation.

### Immunofluorescence

Samples were rinsed with PBS-/-, fixed with 4% paraformaldehyde for 20-30 minutes, then rinsed again with PBS-/-. Samples were blocked and permeabilized with 3% Donkey Serum (Jackson Immuno, Catalog number NC9624464) and 0.1% Triton X-100 in PBS-/-(blocking buffer) for 30 minutes, then incubated with designated primary antibody dilutions in blocking buffer overnight at 4°C. The following day samples were washed three times in 0.1% Tween-20 (PBST) for 5-20 mins per wash, then incubated with designated secondary antibody dilutions for 30 minutes to 1 hour covered in foil to minimize light exposure. Then samples were washed three times for 5-20 mins per wash in PBST. Samples were imaged and stored in PBS. All steps were carried out at room temperature unless otherwise specified.

| Antibodies |  |  |
| --- | --- | --- |
| Anti-mouse GATA3 antibody | Thermo Fisher Scientific | Cat #: MA1-028<br>RRID: AB_2536713 |
| Anti-goat Brachyury antibody | R & D Systems | Cat #: AF2085<br>RRID: AB_2200235 |
| Anti-rabbit SOX2 antibody | Cell Signaling Technology | Cat #: 3579<br>RRID: AB_2195767 |
| Alexa Fluor™ 568 Donkey anti-Mouse | Thermo Fisher Scientific | Cat #: A10037<br>RRID: AB_11180865 |
| Alexa Fluor™ 647 Donkey anti-Goat | Thermo Fisher Scientific | Cat #: A-21447<br>RRID: AB_2535864 |
| Alexa Fluor™ 488 Donkey anti-Rabbit | Thermo Fisher Scientific | Cat #: A-21206<br>RRID: AB_2535792 |
| Alexa Fluor™ 405 Donkey anti-mouse | Thermo Fisher Scientific | Cat #: A-48257<br>RRID: AB_2884884 |

### Single cell RNAseq sample collection and processing

First, 384 well plates were prepared for sample collection and processing. Vapor-Lock (QIAGEN, 981611) was dispensed into each well of a 384-well plate (Bio-Rad, HSP3801) using a 12-channel pipette. All downstream dispensing into 384-well plates was performed using a Nanodrop II liquid handling robot (BioNex Solutions). Uniquely barcoded reverse transcription (RT) primers containing a 6-nucleotide UMI, lysis buffer (0.175% IGEPAL CA-630, 1.75 mM dNTPs, 1:1,250,000 ERCC RNA spike-in mix (Ambion, 4456740), and 0.19 U RNase inhibitor (Clontech, 2313A)) was added to each well. The RT primers used here were described by Grün et al^73^.

Single-cell suspensions were made by dissociating cells using Accutase, inactivating using mTeSR Plus, then washing and resuspending in 1 ml of cold 1x DPBS (Gibco, 14190144) before being passed through a 35 µm nylon mesh cell strainer into a 5 mL round bottom polystyrene tube (Corning, 352235) that was kept on ice. Single cells were FACS sorted into individual wells of a 384-well plate and immediately placed over ice.

After sorting, cells were first lysed, then RT was conducted by adding reverse transcription mix (0.7 U RNase OUT (Invitrogen, 10777-019), 2.34x first strand buffer (Invitrogen, 10777-019), 23.34 mM DTT (Invitrogen, 10777-019), 3.5 U Superscript II (Invitrogen, 18064-071)) to each well, and the plates were incubated at 42°C for 1 hour and 15 minutes, 4°C for 5 minutes, 70°C for 10 minutes. Next, second strand synthesis was carried out by adding second strand synthesis mix (1.74x second strand buffer (Invitrogen, 10812-014), 0.35 mM dNTP (Invitrogen, 10812-014), 0.14 U *E. coli* DNA Ligase (Invitrogen, 18052-019), 0.56 U *E. coli* DNA Polymerase I (Invitrogen, 18010-025), 0.03 U RNase H (Invitrogen 18021-071)) to each well and incubating at 16°C for 2 hours. 1 µL of nuclease-free water was added into each well to minimize losses during downstream pooling. The endogenous mRNA molecules in a well now all contain the same RNA capture barcode, indicating a unique cell, which allows for pooling. Reaction wells receiving different barcodes were pooled using a multichannel pipette, and the oil phase was discarded.

1.0x AMPure XP paramagnetic solid phase reversible immobilization (SPRI) size selection beads (Beckman Coulter, A63881) were used to further purify samples. Samples were eluted in 30 µL nuclease-free water and concentrated using an Eppendorf Vacufuge plus. Linear amplification of molecules via in vitro transcription (IVT) was performed using the MEGAscript T7 transcription kit (Ambion, AMB13345). Samples were then incubated at 37°C with the lid set at 70°C for 13 hours to produce aRNA (amplified RNA). ExoSAP-IT treatment of samples was performed to remove unwanted primers. RNA fragmentation was conducted next using 0.25x fragmentation buffer (200 mM Tris-acetate, pH 8.1, 500 mM KOAc, 150 mM MgOAc) followed by an incubation at 94°C. Immediately after incubation, samples were moved to ice and 0.1x stop buffer (0.5 M EDTA) was added. Sample volumes were increased to 50 µL by adding 19.75 µL of nuclease-free water.

Next, an RNA sample cleanup was conducted by adding 0.825x RNAClean XP beads (Agencourt, A64987) to samples. The samples + beads mixture were placed on a magnetic stand, and beads were washed twice using 80% ethanol then allowed to air dry before samples were eluted in nuclease-free water. Another RT reaction was conducted next with a random hexamer primer that contains an Illumina adaptor handle. Random hexamer primers (20 µM) and dNTP mix were added to 5 µL of the previously eluted aRNA, incubated for 5 minutes at 65°C and then placed over ice. RT reaction mixture (first strand buffer, DTT, RNaseOUT, and SuperScript II enzyme) was added to the aRNA samples then incubated in a thermal cycler for 10 minutes at 25°C and 60 minutes at 42°C. A final PCR amplification was conducted next to generate full length Illumina libraries. A uniquely indexed RNA RPIX primer (10 µM), RNA RP1 primer (10 µM), nuclease-free water, and PCR mix (NEB, M0541S) was added to half the sample volume. Samples were then amplified in a thermal cycler for 30 seconds at 98°C, and 6-12 cycles of: 10 seconds at 98°C, 30 seconds at 60°C, 30 seconds at 72°C, followed by 10 minutes at 72°C, and finally holding temperature at 4°C.

To purify complete Illumina library samples, 0.8X AMPure XP SPRI size selection beads were used and washed again as described above. After the beads were allowed to air dry, 50 µL of nuclease-free water was added. Beads were resuspended again, incubated at room temperature for 5 minutes, and then placed on the magnetic stand for 5 minutes. 50 µL of the sample supernatant was then transferred to a new tube. This purification process was then repeated (0.8x bead and two 80% ethanol washes), and beads were allowed to air dry (∼5 minutes). The final product was eluted in 20 µL of nuclease-free water. Library fragment size distribution was determined using a 2100 Bioanalyzer.

### Single cell RNAseq data processing and gene expression analysis

Sequencing FASTQ data files were processed using custom Perl scripts on a Linux high performance computing (HPC) cluster at the University of California, Santa Barbara. Sequencing data were post-processed and analyzed using Seurat (version 5.3.0), R (version 4.4.2), and RStudio (version 2025.05.0+496).

Paired-end reads obtained by CEL-Seq2^74^ were aligned to the transcriptome using STAR aligner (version 2.7.8a) to the RefSeq gene model based on the human genome release hg38 incorporating a collection of 92 ERCC spike-in molecules. In cases where a read aligned to multiple locations, it was evenly distributed across those locations. Gene isoforms were consolidated into a single gene count, and the unique molecular identifiers (UMIs) were used to remove duplicate reads and generate transcript counts at the single-molecule level for each gene within individual cells. Genes that were not detected in at least one cell were excluded from subsequent analyses.

Single-cell RNA-seq data were processed and analyzed using R and the Seurat package following standard workflows. Quality control filtering was applied to remove low-quality cells and potential multiplets, where cells were retained if they expressed more than 1000 unique genes, resulting in a total of 1299 cells for analysis. Following filtering, gene expression counts were normalized using the log-normalization function, NormalizeData and scaled using the ScaleData function. Highly variable features were identified using FindVariableFeatures with default parameters (selection.method = “vst”, nfeatures = 2000).

Dimensionality reduction was performed using principal component analysis (RunPCA ()), and the first 10 principal components were used for clustering. A shared nearest neighbors (SNN) graph was constructed with FindNeighbors, and clusters were identified using the Louvain algorithm via FindClusters with a resolution of 0.5 unless otherwise noted. Uniform Manifold Approximation and Projection (UMAP) was used to visualize cells in two dimensions (RunUMAP).

To account for potential batch effects across samples (e.g., plates, libraries, or conditions), data integration was performed using Harmony. PCA embeddings were corrected using IntegrateLayers with method = HarmonyIntegration and orig.reduction = “pca”. The resulting batch-corrected embeddings (integrated.harmony) were used to construct a shared nearest neighbor graph using FindNeighbors, identify clusters (FindClusters, resolution = 0.5), and generate a two-dimensional UMAP embedding (RunUMAP, reduction = “integrated.harmony”). Clustering results were annotated using both metadata (e.g., sample origin, experimental condition) and known marker genes (**see supplementary note 1**). Differentially expressed genes for each cluster were identified with FindAllMarkers using the Wilcoxon rank-sum test, filtering for genes with an adjusted p-value < 0.05 and log2 fold change > 1. For visualization, heatmaps, violin plots, feature, and dot plots were generated using DoHeatmap, VlnPlot, FeaturePlot, and DotPlot, respectively.

### Imaging

All micropatterned fixed cell images shown were imaged using a Nikon W2 SoRa spinning-disk confocal microscope. Un-patterned critical window experiments were imaged using the CellVoyager CQ1 Benchtop High-Content Analysis System (Yokogawa Electric Corporation). Four channels were imaged corresponding to DAPI, Alexa488, and Alexa647 conjugated antibodies, and endogenous b-catenin tdmruby. Images were exported as .tiff files and analyzed using Fiji, and custom MATLAB and Python software, enabling quantification of the spatial distribution and intensity of cell fate markers across various light treatments. Live cell confocal imaging was performed using a Nikon W2 SoRa spinning-disk confocal microscope equipped with incubation chamber maintaining cells at 37°C and 5% CO2.

#### Image Analysis

##### Marker Quantification

We developed in-house MATLAB software to extract intensity and positional intensity of imaged colonies. The quantification of each imaged colony was performed in MATLAB using the same general workflow: background subtraction > dilation > nuclei detection > measurement. Subcellular segmentation of nuclear fluorescence was performed using DAPI brightness, size, and circularity to mask nuclei. Mean fluorescent intensity of ROIs were measured and subsequently processed. It is described in detail by Rufo et al^13^.

##### Quantifying WNT signaling (Python)

To quantify active WNT signaling, we developed a custom image processing algorithm to differentiate the active, non-membrane β-catenin from the inactive, membrane-bound pool. The algorithm was implemented using Python with the OpenCV and scikit-image libraries for image processing, along with numpy for numerical calculations and is described in more detail by Rufo et al ^12^.

##### Pattern Fidelity Index Metric

In the 2D gastruloid, Brachyury patterns in a ring that is a few cell widths in from the edge of the colony. The region of maximum BRA intensity is in this ring, and the BRA signal in the center of the colony should be much lower (**fig S4A**). We noticed some of our light treatments recapitulated this trend (with BRA intensity higher close to the edge, and lower in the center), while others resulted in BRA signal that was evenly distributed across the colony. Even further, we observed light treatments that resulted in higher BRA intensity in the center of the colony, and lower on the edge representing an inverted patterning phenotype (BRA germ layer in the center of the colony, and SOX2+ surrounding). Therefore, we developed a metric that would quantitatively capture how faithfully the light treatment recapitulated the expected radial symmetry of the 2D gastruloid. First, we determined the 3 bin regions of max and min intensity in our light treatments (**fig S4B**) and selected Bins 1 - 4 (11 µm - 40 µm) as the representation of the max intensity and bins 21-24 (211-240 µm) which is the center of the colony. Then we quantified the PFI for each light treatment based on the BRA intensity in the edge and center regions quantified in the BRA radial profiles.

The PFI is calculated as follows:

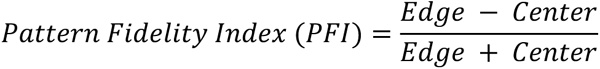

This metric is bounded and easily interpretable for our high throughput data. A PFI value that is negative (less than -0.1) represents inverted patterning, a value close to 0 (between -0.1 and 0.1) represents uniform BRA patterning, and the larger the value, the greater degree of pattern fidelity (more separation/ greater difference between the brachyury intensity at the edge compared to the center). Notably, BMP4 patterned colonies achieve a PFI value of greater than 0.32 while low values (0.1–0.32) indicate less radial symmetry.

We use two different edge regions to calculate the PFI depending on the context (the edge region used is explicitly stated in the main text). The 3-bin max of a normal BMP4 patterned colony is Bins 6-8 (60-80 µm). Optogenetically patterned colonies in the full temporal screen (**Figures 2 and 3**) result in a different BRA ring region because the outer extraembryonic ring is not clearly defined (**Fig 5A, fig S2C**) likely due to the uniform activation of WNT, resulting in BRA signal that reaches to the colony edge. Therefore, to highlight the differences between optogenetically patterned colonies from the temporal screen (**Figs 2 and 3**), we use the 3-bin max region identified for the screen (Bins 1-4 or 11-40µm) which is notably closer to the edge than in a normal BMP4 patterned colony. To highlight full rescue of germ layer patterning (Figure 5) we quantified the PFI using the expected target region of max intensity for a normal BMP4 patterned colony.

The pattern fidelity index calculation for full optical rescue gastruloids stained for GATA3, BRA and SOX2 (**Fig 5G**) was quantified manually since no channel was available for the nuclear stain. Briefly, ROIs corresponding to the 3-Bin “Edge” and “Center” regions were identified in Fiji, and the brachyury pixel intensity was quantified for each region. To determine the “Edge” ring pixel intensity, the integrated density (pixel intensity * ROI area) of a circular ROI 51µm from the edge of the colony was subtracted from the integrated density of a circular ROI 80 µm from the edge of the colony. This value was converted back to mean intensity for the PFI calculation.

## Data availability

Sequencing data have been deposited in the Gene Expression Omnibus (GEO) database accession code GEO: GSEXXXXXX (Data can be accessed using the following token: XXXXXXXXXXXXXXX).

## Notes

### Competing Interest Statement

Maxwell Z Wilson is co-founder and CSO of Integrated Biosciences.

